# Immunodominant T-cell epitopes from the SARS-CoV-2 spike antigen reveal robust pre-existing T-cell immunity in unexposed individuals

**DOI:** 10.1101/2020.11.03.367375

**Authors:** Swapnil Mahajan, Vasumathi Kode, Keshav Bhojak, Coral M. Magdalene, Kayla Lee, Malini Manoharan, Athulya Ramesh, HV Sudheendra, Ankita Srivastava, Rekha Sathian, Tahira Khan, Prasanna Kumar, Papia Chakraborty, Amitabha Chaudhuri

**Affiliations:** MedGenome, Bangalore, India; MedGenome, Foster City, USA

**Author notes:** Equal contribution (Swapnil Mahajan, Vasumathi Kode, Keshav Bhojak, Coral M. Magdalene).

## Abstract

The COVID-19 pandemic has revealed a range of disease phenotypes in infected patients with asymptomatic, mild or severe clinical outcomes, but the mechanisms that determine such variable outcomes remain unresolved. In this study, we identified immunodominant CD8 T-cell epitopes in the RBD and the non-RBD domain of the spike antigen using a novel TCR-binding algorithm. A selected pool of 11 predicted epitopes induced robust T-cell activation in unexposed donors demonstrating pre-existing CD4 and CD8 T-cell immunity to SARS-CoV-2 antigen. The T-cell reactivity to the predicted epitopes was higher than the Spike-S1 and S2 peptide pools containing 157 and 158 peptides both in unexposed donors and in convalescent patients suggesting that strong T-cell epitopes are likely to be missed when larger peptide pools are used in assays. A key finding of our study is that pre-existing T-cell immunity to SARS-CoV-2 is contributed by TCRs that recognize common viral antigens such as Influenza and CMV, even though the viral epitopes lack sequence identity to the SARS-CoV-2 epitopes. This finding is in contrast to multiple published studies in which pre-existing T-cell immunity is suggested to arise from shared epitopes between SARS-CoV-2 and other common cold-causing coronaviruses. Whether the presence of pre-existing T-cell immunity provides protection against COVID-19 or contributes to severe disease phenotype remains to be determined in a larger cohort. However, our findings raise the expectation that a significant majority of the global population is likely to have SARS-CoV-2 reactive T-cells because of prior exposure to flu and CMV viruses, in addition to common cold-causing coronaviruses.

## INTRODUCTION

Uncovering the immunological responses to COVID-19 infection will help in designing and developing next-generation therapies and manage the treatment of critical COVID-19 patients. Many host factors associated with mild or severe disease symptoms have been reported. For example, leukopenia, exhausted CD8 T-cells, higher levels of T_H_2 cytokines in serum, a high titer of neutralizing antibodies, blunted interferon response, dysregulation of the myeloid cell compartment, activated NK cells, and the size of the naïve T-cell compartment is associated with critically ill patients (1-4). This wide range of variable factors shares a common immunological underpinning – that of a systemic dysregulation in immune homeostasis due to the failure of the host immune system to clear the virus during the early stages of the infection (5). Many studies have shown that clearance of respiratory viruses requires CD8 T-cell immunity (6). A delay in the activation of CD8 T-cells and a lack of early IFN-γ production lead to an increase in viral load triggering overactivation of the innate and the adaptive arm of the immune system leading to a loss of immune homeostasis resulting in severe disease phenotype, including death. Therefore, an early wave of strong CD8 T-cell response may delay viral titer build-up, allowing rapid clearance of the virus by the immune system without perturbing immune homeostasis.

Healthy humans not exposed to COVID-19 show pre-existing CD4 and CD8 T-cell immunity to SARS-CoV-2 antigens (7-9). The pre-existing immunity to CD4 and CD8 T-cells was detected against structural and non-structural SARS-CoV-2 proteins by overlapping 15-mer peptide pools. The existence of a pool of SARS-CoV-2-reactive T-cells in unexposed individuals is thought to arise from coronaviruses that cause common cold (8, 10). Whether pre-existing immunity provides any protection to SARS-CoV-2 infection, or contribute to a faster recovery from infection remains speculative. Besides, it is unclear whether a pre-existing immunity, involving either CD4 or CD8 T-cells, or both, is required for maximal protection. Identifying robust pre-existing immunity against SARS-CoV-2 in the healthy population can be used as a measure to assess the mode of recovery and also viral spread in the global population.

In this study, we identified strong CD8 T-cell-activating epitopes from SARS-CoV-2 spike protein by a combination of epitope prediction and T-cell activation assays in healthy donors unexposed to SARS-CoV-2. The rationale for identifying epitopes that favor CD8 T-cell activation was two-fold. First, robust CD8 T-cell activating epitopes can be formulated as second-generation vaccines for short and long-term protection against viral infection. Second, detection of pre-existing immunity in healthy donors using epitopes that favor CD8 T-cell activation may provide a framework to understand the complex immune responses observed in clinical settings. It may also shed light on the differences in morbidity and mortality in different population groups across the globe.

We developed a proprietary algorithm OncoPeptVAC to predict CD8 T-cell activating epitopes across the SARS-CoV-2 proteome. OncoPeptVAC predicts binding of the HLA-peptide complex to the T-cell receptor (TCR). We selected a cocktail of eleven 15-mer peptides with a broad class-I and class-II coverage and favorable TCR engagement predicted by the algorithm. The cocktail of peptides was tested for T-cell activation in healthy donors from the USA and India unexposed to COVID-19. We observed higher CD8 T-cell activation by the 11-peptide pool compared to the overlapping 15-mer peptide pools from the spike-S1 and S2 proteins. Homology analysis of the selected peptides with other coronavirus spike proteins indicated a lack of significant amino acid identity with any of the 11 peptides, suggesting engagement of one or more peptides in the pool to cross-reactive TCRs from other viruses, not particularly from a coronavirus. Bulk and single-cell TCR analysis revealed expanded clonotypes recognizing epitopes from CMV, Influenza-A, and other viruses to which most of us are exposed. Taken together, our findings support that strong pre-existing CD8 T-cell immunity in unexposed donors is contributed by cross-reactive TCRs from other viruses. Significantly, we discovered multiple immunodominant epitopes in our predicted pool of peptides that favored CD8 T-cell activation. Finally, we show that our cocktail of 11-peptides induced a robust immune response in convalescent patients demonstrating that these peptides are recognized by infected patients.

Taken together, our study uncovered strong pre-existing CD8 T-cell immunity against SARS-CoV-2 using a small set of 11 epitopes that engaged cross-reactive TCRs recognizing epitopes from other viruses, not necessarily common cold viruses belonging to the coronavirus family as hypothesized by other studies. Additionally, our findings provide a basis for the generation of herd immunity against COVID-19 without prior SARS-CoV-2 infection.

## RESULTS

### Prediction of immunogenic epitopes favoring CD8 T-cell activation

A deep CNN model OncoPeptVAC was implemented to predict the immunogenicity of the peptide-based only on the peptide and HLA sequences. A total of 8,870 immunogenic and non-immunogenic peptide-HLA pairs were obtained from the IEDB (11). The BLOSUM encoding was used to represent the peptide and HLA molecules. The BLOSUM substitution scores encode evolutionary and physicochemical properties of the amino acids (12). In addition, hydrophobicity indices and predicted HLA binding scores were also used to represent the peptide and HLA sequences.

OncoPeptVAC used the CNN model with multiple 2D convolutional layers combined with max-pooling to confirm the additive effect of different input features on the performance of the model. All the model versions were trained using 5-fold cross-validation. The AUCs of the final model was 0.87 based on a blind test dataset (Figure 1A). The prediction algorithm showed a sensitivity of 0.64 and a specificity of 0.84 based on the score cut-off of 0.2 (Figure 1B). By increasing the cut-off score to 0.5, the specificity could be further increased to 96.8 with a concomitant loss in sensitivity. OncoPeptVAC reduced the number of false positives significantly compared to the HLA-binding rank (Compare Figures 1B and C) reducing the number of epitopes that needed to be screened in a T-cell activation assay by 30% to identify true immunogenic epitopes. For example, to identify 50% (119 out of 238) of the immunogenic peptide-HLA pairs present in the blind test dataset, 256 top peptide-HLA pairs from OncoPeptVAC prediction needed to be screened, compared to 753 top peptide-HLA pairs predicted by netMHCpan-4.0.

**Figure 1.**
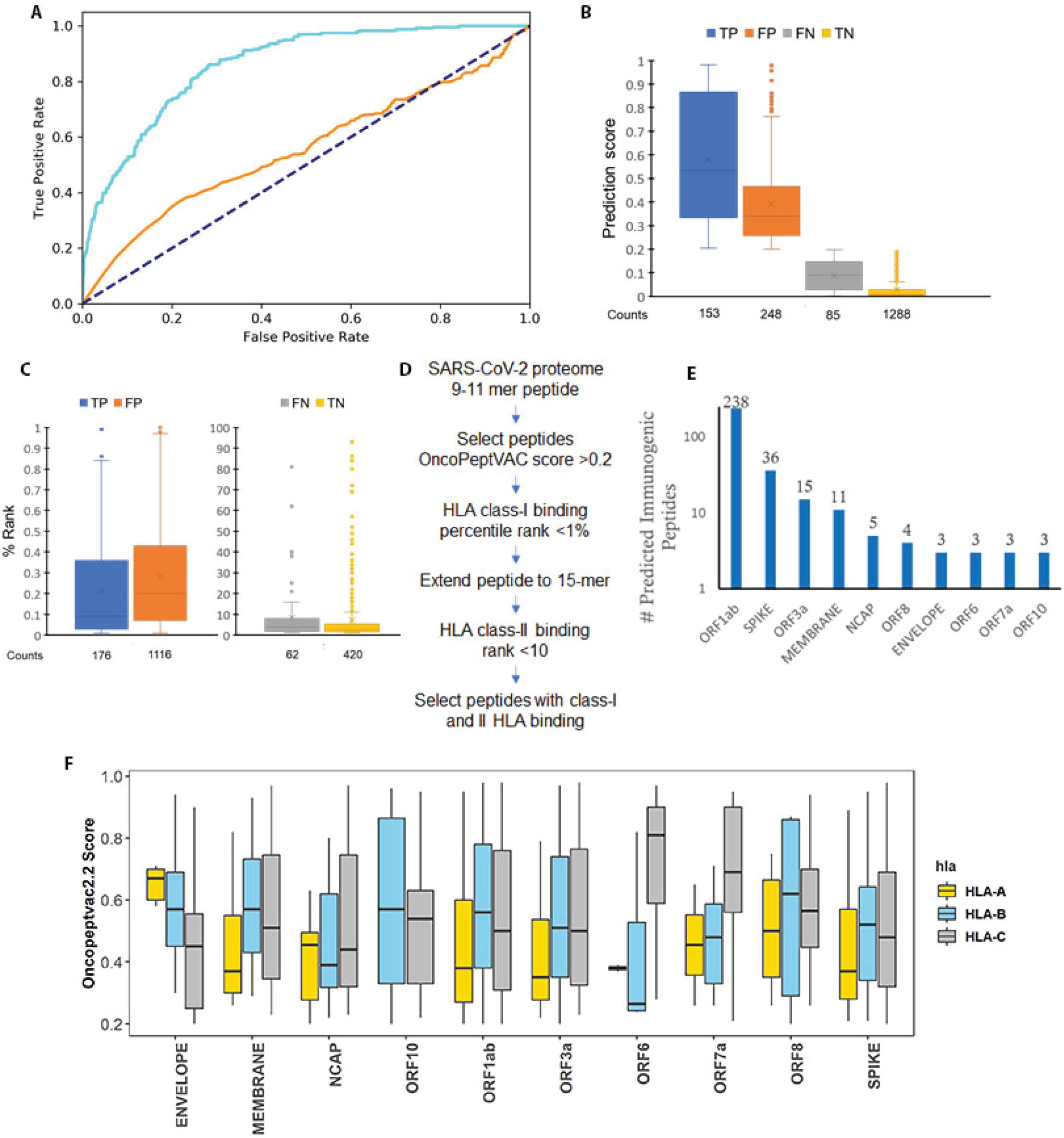
Identification of immunogenic epitopes from SARS-CoV-2 by OncoPeptVAC. **A**. ROC curves of OncoPeptVAC TCR-binding and netMHCpan-4.1 HLA binding algorithms. A blind dataset of non-immunogenic or immunogenic HLA class-I binding T-cell epitopes from IEDB was used to assess the performance of OncoPeptVAC (cyan). The HLA-binding affinity of the epitopes expressed as percentile rank <1% was used to assess the performance of netMHCpan-4.1 in predicting true immunogenic epitopes (orange). **B**. Separation of immunogenic from non-immunogenic epitopes by OncoPeptVAC score. **C**. Separation of immunogenic from non-immunogenic epitopes by HLA-binding percentile rank. **D**. Schematic showing the steps used to identify immunogenic epitopes from SARS-CoV-2 proteome. **E**. Number of immunogenic epitopes identified in different SARS-CoV-2 antigens. **F**. HLA-A, B and C-restricted epitopes from SARS-CoV-2 proteome.

The prediction algorithm was applied to the SARS-CoV-2 proteome and screened against 23 class-I HLAs covering over 98% of the world population. A schematic of the *in silico* screening approach is shown in Figure 1D. Briefly, 9-11-mer peptides from the SARS-CoV-2 proteome were screened for TCR-binding against 23 HLA, and peptides with OncoPeptVAC score >0.2 were analyzed for class-I HLA binding. Peptide-HLA pairs with a high predicted binding affinity (<1 percentile rank) were selected, their length extended to 15-mer and screened for class-II HLA binding. Peptides with favorable TCR binding and class-I/II HLA binding features were selected for further validation. The number of predicted immunogenic epitopes from SARS-CoV-2 protein-coding genes is shown in Figure 1E. The distribution of OncoPeptVAC scores against different class-I HLA genes indicates a higher number of favorable TCR-binding peptides for HLA-B and C compared to HLA-A (Figure 1F). Natural biases in HLA-restrictions have been reported for immunogenic HIV epitopes (13).

### T-cells from unexposed donors respond to OncoPeptVAC-predicted peptides

We performed T-cell activation assay using the selected 11 epitopes from the SARS-CoV-2 spike antigen in unexposed donors. The 15-mer peptides are distributed across different segments of the RBD and the non-RBD regions of the spike antigens and few peptides carry ACE2 receptor binding sites (Figure 2A and Table-1). Many predicted peptides reside in flexible regions of the spike protein that could favor efficient processing and presentation.

**Table-1.** Peptides selected by OncoPeptVAC and used in T-cell activation assays. The OncoPeptVAC score was calculated for each HLA-peptide pair (see Methods). The HLA-peptide binding affinity percentile rank was calculated by netMHCpan4.0.Peptides. Peptides having >0.2 OncoPeptVAC score, <1 class-I HLA-binding percentile rank, and ≤10 class-II HLA binding percentile rank were selected.

| Peptide ID | # Class-I HLA restriction | #Class-II HLA restriction | OncoPeptVAC score for individual HLAs (range) | Class-I binding percentile rank (range) | Class-II binding percentile rank (range) | Presence in spike peptide pools | Other features |
| --- | --- | --- | --- | --- | --- | --- | --- |
| Pep-1 | 3 | 0 | 0.21-0.51 | 0.27-0.77 | NA | Yes | One contact residue for neutralizing Ab |
| Pep-2 | 0 | 2 | NA | NA | 7.6-7.8 | No | 4 ACE contact residues and 2 contact residues for neutralizing Ab |
| Pep-3 | 1 | 3 | 0.27 | 0.83 | 1.5-8.5 | No |  |
| Pep-4 | 7 | 9 | 0.43-0.94 | 0.02-0.25 | 0.58-10.0 | No |  |
| Pep-5 | 2 | 1 | 0.24-0.28 | 0.45-0.49 | 3.1 | Yes | 4 ACE contact residues |
| Pep-6 | 6 | 10 | 0.28-0.9 | 0.37-0.97 | 0.27-9.6 | No |  |
| Pep-7 | 11 | 18 | 0.21-0.94 | 0.06-0.8 | 0.49-9.3 | Yes |  |
| Pep-8 | 14 | 16 | 0.21-0.7 | 0.01-0.87 | 0.97-10.0 | No |  |
| Pep-9 | 27 | 13 | 0.22-0.96 | 0.02-0.89 | 1.8-8.8 | No |  |
| Pep-10 | 4 | 9 | 0.23-0.83 | 0.19-0.71 | 0.45-8.8 | No | 5 ACE contact residues and 2 contact residues for neutralizing Ab |
| Pep-11 | 6 | 2 | 0.22-0.88 | 0.44-0.9 | 7.2-8.0 | No |  |

**Figure 2.**
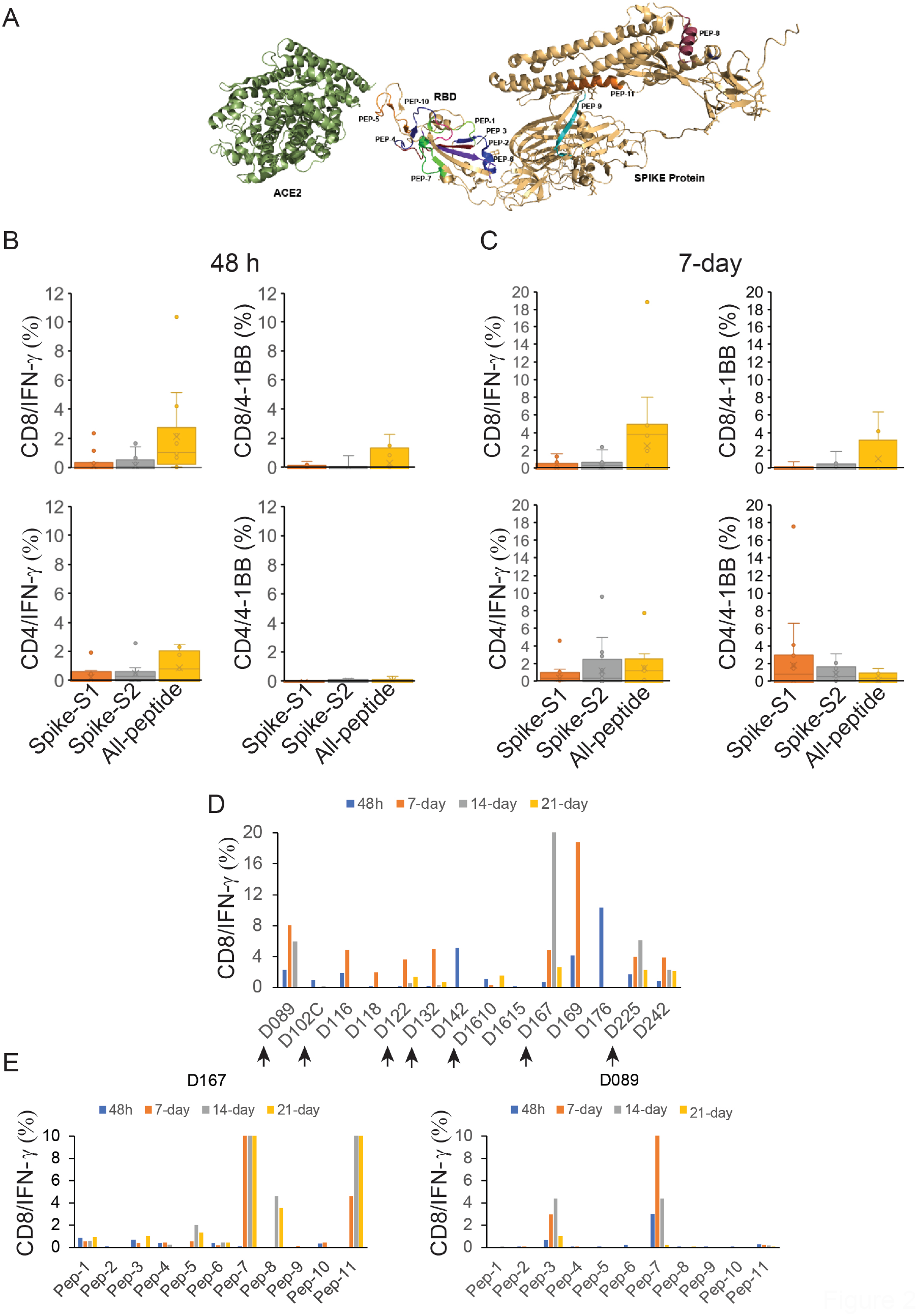
T-cell reactivity by SARS-CoV-2 Spike peptides. Reactivity was determined by intracellular IFN-γ staining and surface expression of 4-1BB by FACS after stimulation of PBMCs from unexposed donors (n=14) using separate pools of Spike-S1 and S2 peptides and the 11-Peptide-mix predicted by OncoPeptVAC. **A**. Structure of Spike – ACE2 receptor complex showing the location of the 11-peptides predicted by OncoPeptVAC. **B**. T-cell activation after 48h incubation with the peptides. **C**. T-cell activation after 7-day incubation with the peptides. **D**. Kinetics and magnitude of CD8 T-cell activation in unexposed donor PBMCs. **E-F**. CD8 T-cell response of donors D167 and D089 to individual peptides from the 11-peptide-mix.

We screened PBMCs from 17 unexposed donors from the US collected between 2016 – 2018 and India (2015 – 2017), much before SARS-CoV-2 was recognized as a global pandemic (Table-S1). Activation of T-cells using the cocktail of 11 peptides (All-peptide) was compared to the responses from Spike-S1 (157 peptides) and S2 (158 peptides) pools (see Methods for assay details). In a 48h assay, 70% of the unexposed donors responded strongly to the predicted peptide mix by inducing intracellular IFNγ ^+^ in both CD4 and CD8 T-cells. The responses to Spike-S1 and S2 peptide pools were weaker (Figure 2B). A strong 48h IFN-γ response suggested recall to pre-existing antigen-experienced CD8 T-cells. The peptide-mix also induced a strong 4-1BB response in CD8 T-cells but not in CD4 T-cells (Figure 2B). Both IFN-γ and 4-1BB levels increased in CD8 T-cells at day-7 by the All-peptide mix compared to the Spike peptide pools (Figure 2C). We observed higher expression of IFN-γ and 4-1BB in the CD4 T-cells by the Spike peptide pools at day-7 suggesting de novo activation (Figure 2C). Although the use of 15-mer peptides is expected to skew the response towards CD4 T-cells, we observed a stronger CD8 T-cell response to the Peptide-mix suggesting that the 15-mer peptide though added exogenously, was processed and presented by class-I HLAs efficiently. Taken together, the results demonstrate that the use of OncoPeptVAC identified potent CD8 T-cell epitopes in the spike antigen that could not have been detected by using large overlapping peptide pools used in T-cell activation assays.

Next, we tested individual peptides from the mix to assess their contribution to T-cell activation. The magnitude and kinetics of IFN-γ induction in CD8 T-cells were variable in different donors (Figure 2D). In most donors, the maximal response was detected by 7-days, but in donors D142 and D176 the response peaked at 48h and declined (Figure 2D). We tested the effect of individual peptides in multiple donors as indicated by the arrows to determine their immunogenicity. As shown in Figure 2E-F and Supplementary Figures S1-S2, multiple peptides from the All-peptide mix induced robust CD8 T-cell IFN-γ and 4-1BB in different donors. The magnitude and the kinetics of response were variable. Peptide-7 was an exceptionally strong CD8 T-cell epitope inducing IFN-γ in 3 out of 7 donors (Figure 2E-F and Supplementary Figure S1). Most immunogenic epitopes activated CD8 T-cells at 48h and achieved >2% activation by 7-day, suggesting that these epitopes engaged pre-existing T-cell immunity in the unexposed donors. In many donors, strong CD4 T-cell response was detected by individual peptides (Figure S3A-J). Interestingly, in many donors, both IFN-γ and 4-1BB expression was induced by multiple individual peptides, although the magnitude of response by the All-peptide mix was not additive (Figure S3A (IFN-γ response and S3F (4-1BB response) confirming that the T-cell activation potential of individual peptides could be masked, when present as a part of a larger peptide pool.

Multiple studies have reported pre-existing T-cell immunity in unexposed donors using spike peptide pools and attributed the response to T-cells recognizing epitopes from common cold-causing coronaviruses to which a large section of the global population is exposed (7, 8, 10). Homology analysis of the selected epitopes (see Methods) indicated that 6 out of the 11 peptides share >67% sequence identity with SARS-CoV and only 1 (Peptide-11) out of the 11 peptides has over 70% identity with multiple coronaviruses (Table 2). Peptide-11 is in the S2 domain of the spike protein and showed ≥1% CD8 T-cell response at 48h in 1 out of 7 donors tested (D167, Figure 1E). However, peptides 3, 6, 7, and 9 lacking significant identity to other coronaviruses (Table 2) showed ≥1% CD8 T-cell activation at 48h in at least one donor out of 7 (Figure 2E-F and Figure S1). Peptide-7 induced high CD8 T-cell activation at 48 h in two donors (D089 and D225 (Figure 2F and Figure S1). Taken together, the data suggest that pre-existing T-cell immunity to these peptides may be derived from cross-reactive TCRs recognizing other viruses.

**Table-2.**
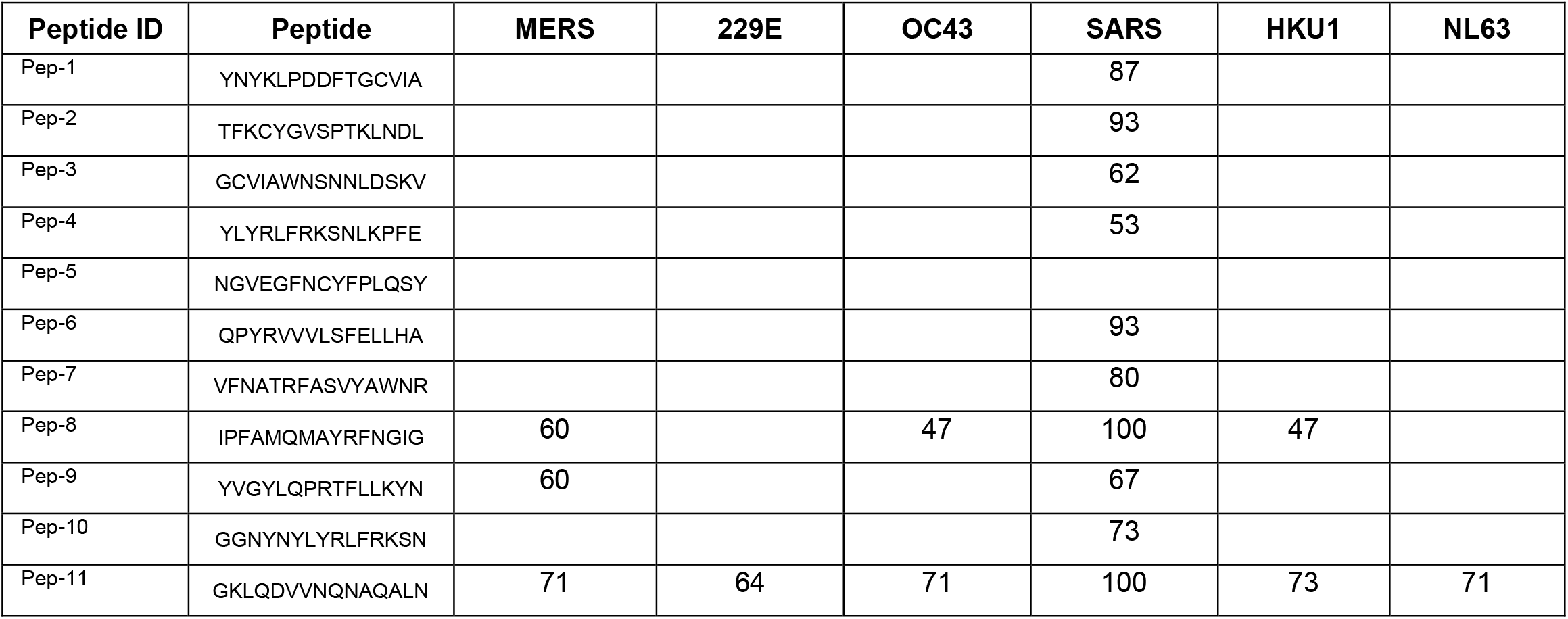
Selected peptides from SARS-CoV-2 and their homology to other coronaviruses. Sequence identity of the shortlisted 15-mer peptides against homologous peptides in other coronaviruses. Peptides were identified based on E-value cut-off of 0.01 and a minimum cut-off length of 11 amino acid residues (see Methods).

### Analysis of antigen-specific CDR3s in responsive donors

To identify CDR3s amplified by individual peptides, or the All-peptide-mix, bulk TCR analysis was performed on antigen-stimulated PBMCs from donors D089 and D225 (see Methods). Both donors showed a robust IFN-γ response to Pep-7, and the Pep-Mix, but not to Pep-1 (Figure S4). Diversity and clonal amplification of unique public and private CDR3s were analyzed at three different time points (Figure 3A-B). Both the donors showed clonal expansion of multiple public CDR3s recognizing HCMV, human herpes virus-5 (HHV-5), and Influenza-A peptides when stimulated with Pep-7 and All-peptide mix, but not with Pep-1 (Table S2). HCMV and HHV-5 CDR3s were expanded in donor D089 (Figure 3A, top panel), whereas D225 showed expansion of HCMV and Influenza-A CDR3s (Figure 3A, bottom panel). Significantly, these CDR3s were not amplified by Spike-S1 and S2 peptide pools or by Pep-1, the latter failed to activate T-cells in these donors. Further, CDR3s recognizing HCMV peptide NLVPMVATV in donors 089 and 225 were different, suggesting that the same antigen engages multiple cross-reactive TCRs in different donors. Next, we analyzed private CDR3s in these two donors to identify novel SARS-CoV-2 antigen-specific CDR3 (Figure 3B). Donor 089 showed a lack of specific amplification of private CDR3s suggesting that the robust CD8 T-cell response detected in this donor may be contributed by the amplified public CDR3s (Figure 3B, top panel). In contrast, two private CDR3s were clonally amplified by Pep-7 and All-peptide mix in D225 suggesting that the T-cell response is derived from both public and non-public TCRs in this donor (Figure 3B, bottom panel). A list of clonally amplified public and private CDR3s detected in the two donors is given in Table S2.

**Figure 3.**
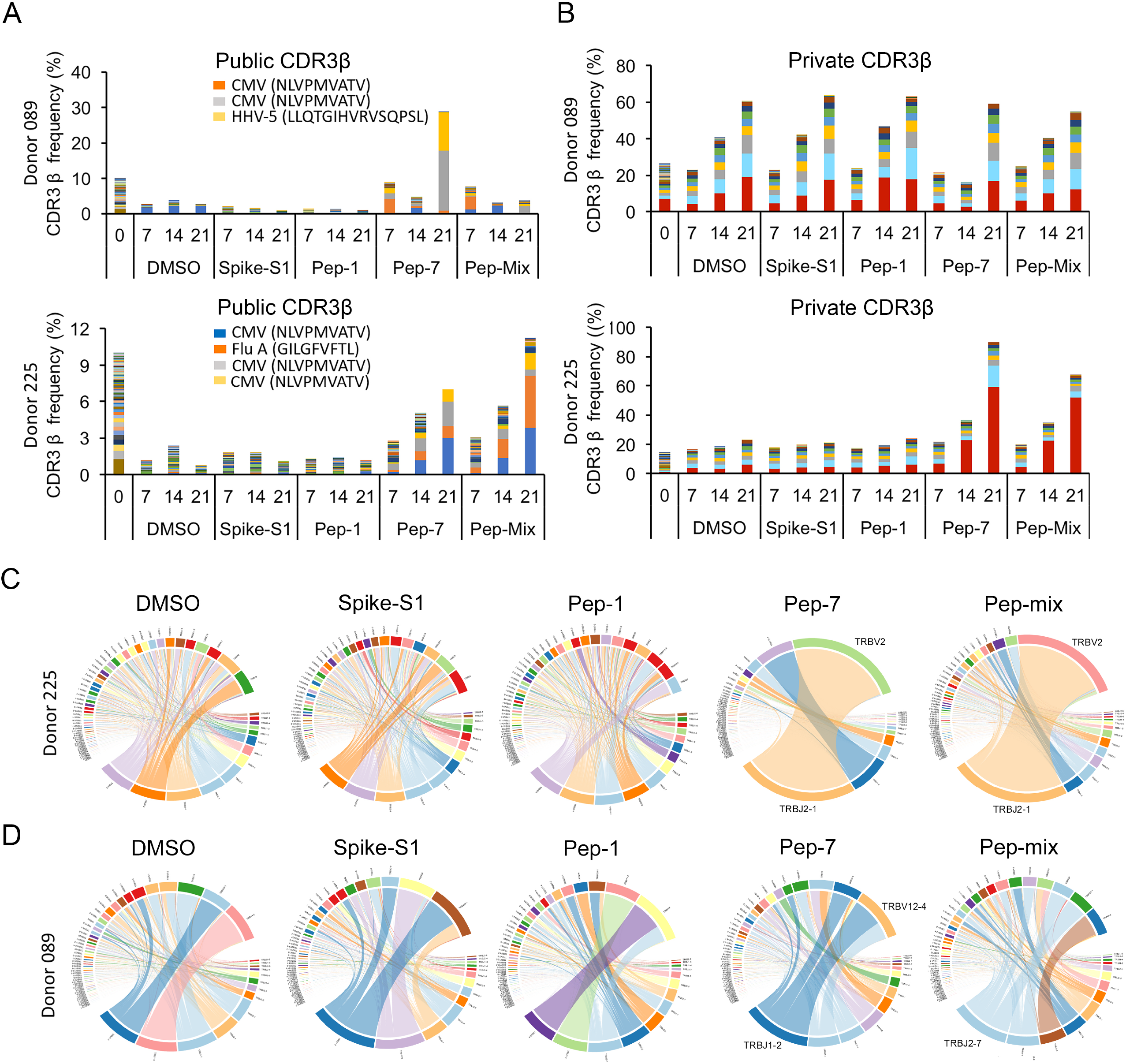
Bulk TCR repertoire analysis after in vitro stimulation of PBMCs at different time points with the indicated peptides in unexposed donors. **A-B**. Expanded public CDR3-βs recognizing shared antigens in the two donors. **C-D**. Expanded private CDR3-βs in the two donors. **E-F**. V-J gene usage in the two donors.

To further investigate the TCR repertoire profile of donors 089 and 225 we analyzed the VDJ gene usage in the bulk CDR3 data. In D225, two V segments TRVB2 and TRVB30 and a J gene TRBJ2-1 were significantly over-represented in Pep7 and Peptide-mix treated samples (Figure 3C), whereas in D089 TRBV12-4 and TRBJ1-2 genes were amplified (Fig. 3D).

### Single-cell transcriptional and TCR profiling of activated T-cells

To characterize the phenotype and functional state of activated T-cells and reveal differences between the different treatments, we performed single-cell sequencing on a 10X platform. Single-cell transcriptomics and TCR data obtained from 3500 – 4500 cells identified 3000 - 3500 unique transcripts (see Methods). Using graph-based clustering of uniform manifold approximation and projection (UMAP) we captured transcriptomes of 4 distinct cell types (Figure 4A and Table S3). Our assay method is enriched for the growth and proliferation of T-cells causing depletion of other immune cell types present in PBMC in a 14-day culture. Three cell types, CD8, γ/δ, and NK-T were detected in all the samples. Compared to DMSO and Pep-7 in which the CD8 T-cell fraction was ∼60%, in spike-S1 and spike-S2 the CD8 T-cell fraction was 50% and 38% respectively. Conversely, the CD4 cluster was expanded in spike-S1 (12%) and S2 (27%) compared to DMSO (7%) suggesting that the spike peptide pools engaged CD4 T-cells (Table S3). The single cell transcriptomic analysis further revealed that Pep-7 induced effector phenotype in the CD8 T-cell cluster by the expression of activation markers IFN-γ, 4-1BB (Figure 4B) TNFRSF9, FAS, and TIGIT (Compare Figure S5D with S5A-C). The top 10 Pep-7-expanded clonotypes were CD27^+^/SELL^-^ suggesting transition towards effector memory phenotype (Figure 4B). Spike-S1 and S2 peptides induced CD27^+^/SELL^-^ T-cells in the CD4 clones 6 and 4 respectively (Figure 4B). Single-cell data revealed amplification of TRBV2 (40%) and TRBJ2-1 (32%) in Pep-7 stimulated T-cells (Figure 4C) confirming the results from the bulk TCR analysis.

**Figure 4.**
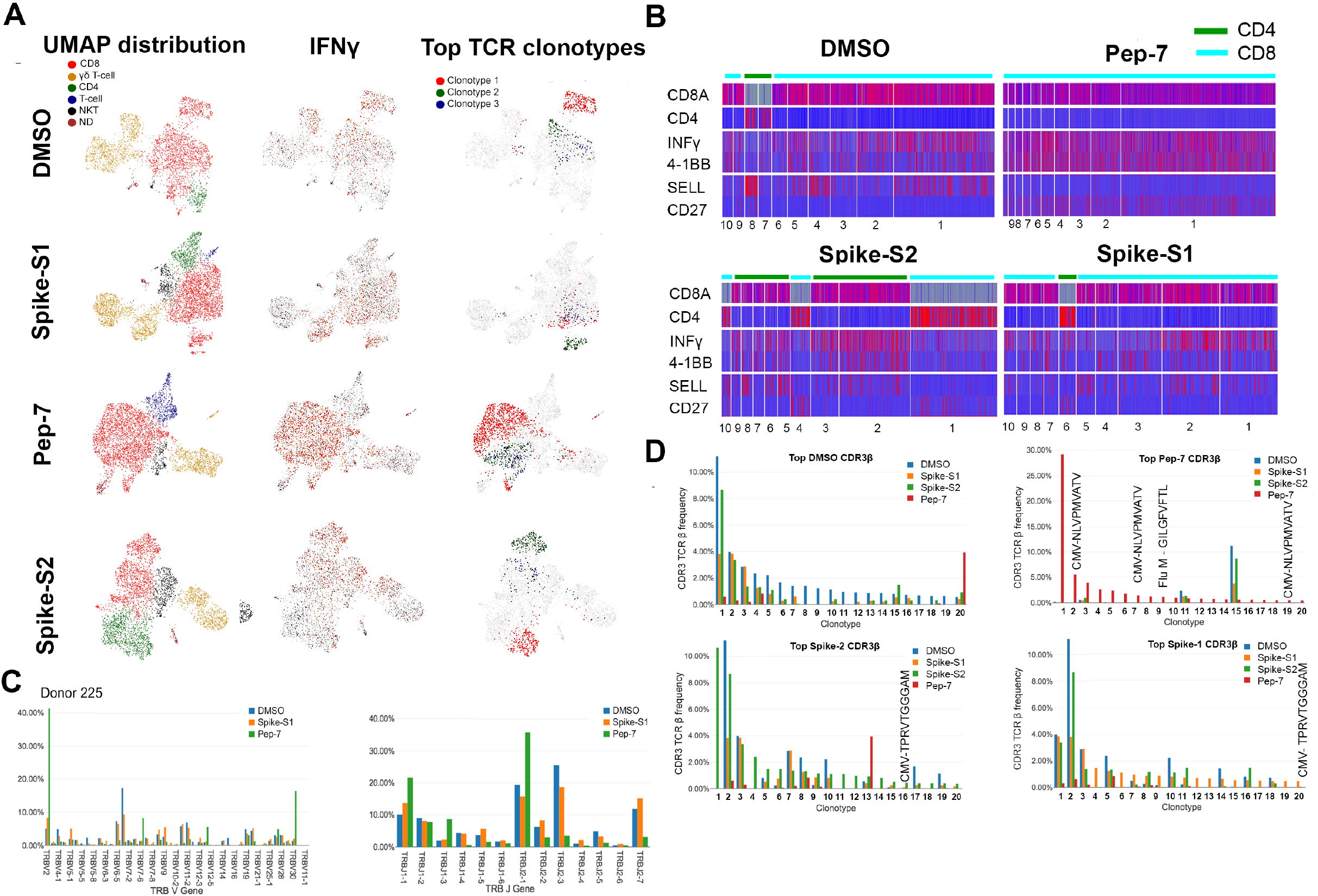
UMAP projection of different cell types identified in unexposed donor D225 after 14 days of in vitro stimulation assay with different antigens. **A**. Clusters of different cell types and their relative proportions present in the assay mixture (left panel). Clusters expressing IFN-γ (middle panel) and the top-3 amplified clonotypes (right panel). **B**. Heat map showing the expression of cell-type and cell-phenotype-specific markers in the top-10 amplified TCR-β clones. **C**. Frequency of CDR3-β recognizing public and private antigens in the top 20 clonotypes. **D**. Amplified V and J-genes in the top-20 clonotypes.

Next, we mapped CDR3-β to specific clones from each treatment (Figure 4D). The DMSO, spike-S1, and S2-treated samples shared many clonotypes among themselves in the same frequency range suggesting weak antigen-induced activation and proliferation of T-cells. However, Pep-7 treated sample was enriched in many CDR3-β clones absent in other samples indicating the specificity of response (Figure 4D, red bars). Four clones among the top-20 clones encoded CMV and flu-specific CDR3-βs in Pep-7 treated sample but not in other samples confirming the findings from the bulk TCR data that the peptide engaged cross-reactive TCRs (Figure 4D). The expanded CDR3-β detected in the Spike-S2 treated sample belonged to CD4 T-cells (Figure 4B, Spike-S2 panel, clonotype-1). Taken together, single-cell TCR analysis demonstrated that the immunogenic SARS-CoV-2 epitope engaged many unique CDR3-βs not shared by spike-1 and spike-2 peptide pools.

The bulk and the single-cell TCR analyses demonstrated that the SARS-CoV-2 epitope identified in this study engaged both cross-reactive public CDR3s and unique CDR3s not associated with known viral antigens and favored the usage-specific V-J genes. Further, the expansion of TRBV-2 and TRBJ2.1 by Pep-7 and by the Pep-mix in D225 confirmed that out of the 11-peptides contained in the Pep-mix, Pep-7 contributed to all of the T-cell responses observed in this donor.

### Antigen-specific clonal expansion and T-cell phenotype

Next, we analyzed the clonal composition and phenotype of T-cells to investigate the dynamics of antigen-specific T-cell response in the treated samples. We analyzed the top-30 clones for their phenotype by the expression of 25 marker genes (Figure S6). In all samples, including DMSO, CD8 T-cell clonotypes were more frequent (Figure S6A-D). As expected, the CD4 T-cell compartment was expanded in Spike-S1 and S2-treated samples (20 and 25% of all clonotypes respectively) compared to DMSO (6.5%) (Figure S6B-C). The CD4 T-cells expressed TNFSF4 (OX-40) suggesting activation, although they failed to express IFN-γ (Figure S6B-C). A few expanded CD4 clones in the Spike-S1 and S2 treated samples showed a high expression of IL17RB suggesting polarization towards a T_H_17 phenotype (Figure S6B-C). In the Pep-7 treated sample, almost all clonotypes in the top-30 were CD8 T-cells. The highly expanded clones expressed multiple T-cell activation markers (Figure S6D). Interestingly, in addition to the activation markers, these cells expressed higher levels of IL2RA (CD25) suggesting differentiation towards an effector memory phenotype (Figure S6D). CD25 expression was low in the CD8 T-cell compartment in other samples. Taken together, the results of the transcriptomic analysis highlighted that the strong immunogenic CD8 T-cell epitope identified in this study preferentially engaged CD8 T-cells pushing them towards an effector and effector memory phenotype. The spike-S1 and S2 peptide pools on the other hand engaged both CD8 and CD4 T-cells and modulated the CD4 T-cells towards a T_H_17 phenotype.

### Response of convalescent COVID-19 patients to predicted epitopes

To assess whether the predicted epitopes are recognized by COVID-19 infected patient T-cells, we tested the All-peptide mix on seven asymptomatic, five with mild-moderate symptoms, and five severe convalescent patients requiring ICU admission (Table S4) and analyzed their CD4 and CD8 T-cell response after 48h. The patients experiencing mild to moderate symptoms exhibited higher induction of IFN-γ in CD8 T-cells (Figure 5A). The IFN-γ induction in CD4 T-cells was higher in the presence of the Spike-S1 peptide pool compared to the All-peptide mix (Figure 5C). Spike-S2 peptide pool induced stronger 4-1BB induction in CD8 and CD4 T-cells compared to the All-peptide mix (Figure 5B-D). Taken together, our results confirm that the epitopes prioritized by the algorithm were recognized by COVID-19 infected patient T-cells, and the IFN-γ response induced by the All-peptide mix was skewed towards CD8 T-cells. Spike peptide pools favored activation of the CD4 compartment in these convalescent subjects in line with our assay results and single cell transcriptome analysis showing a preferential expansion of the CD4 compartment by these peptide pools.

**Figure 5.**
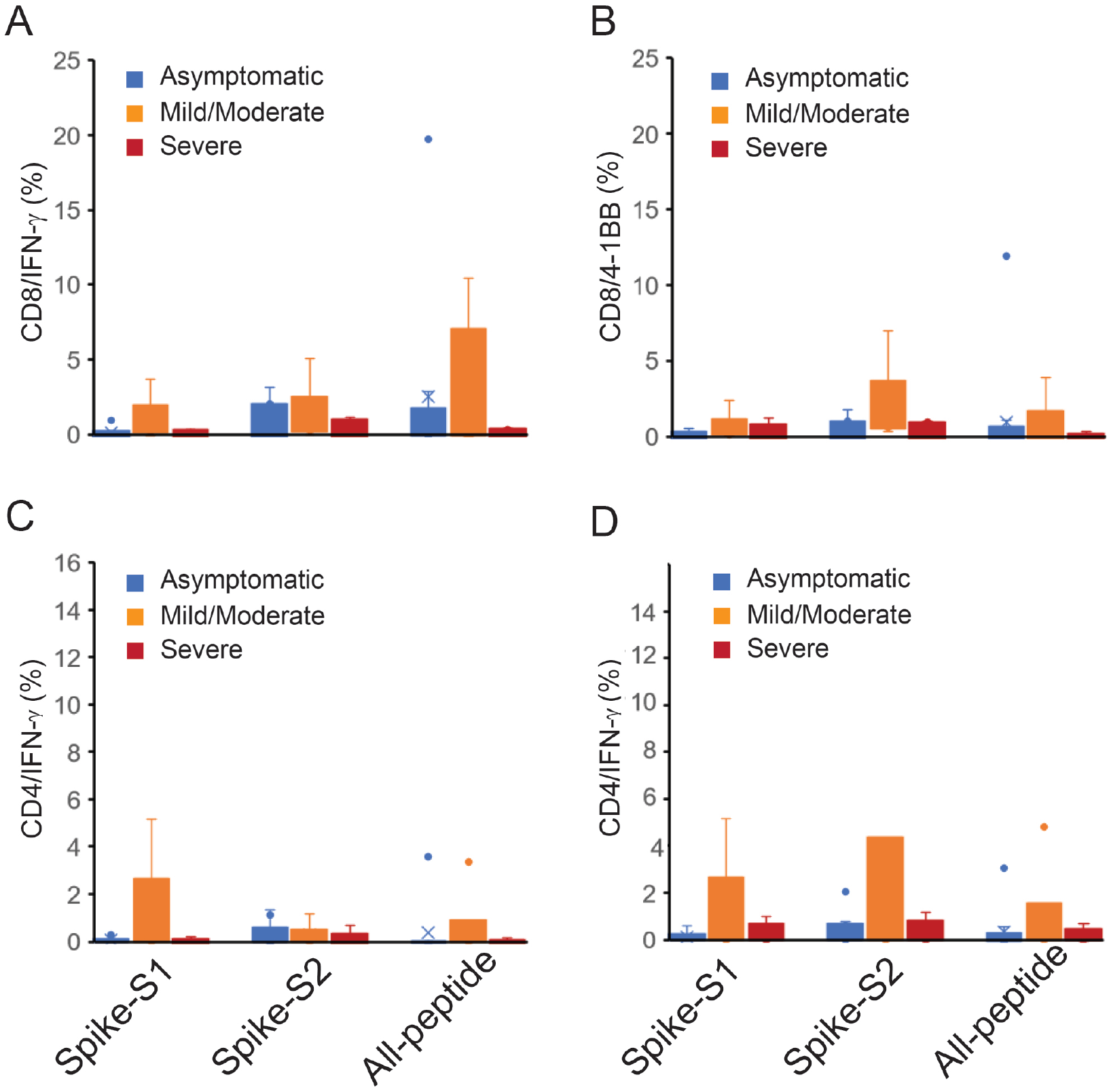
T-cell reactivity to Spike-S1, S2 pools and 11-peptide-mix in asymptomatic, mild-moderate and severe disease after in vitro stimulation for 48 h. **A**. IFN-γ and 4-1BB expression in activated CD8 T-cells. **B**. IFN-γ and 4-1BB expression in activated CD4 T-cells.

## DISCUSSION

A wide array of respiratory viruses induces severe pneumonia, bronchitis, and even death following infection. Despite the immense clinical burden, there is a lack of efficacious vaccines with long-term therapeutic benefit. Most current vaccination strategies employ the generation of broadly neutralizing antibodies, however, the mucosal antibody response to many respiratory viruses is short-lived and declines with age. In contrast, several studies on respiratory viruses have shown the presence of robust virus-specific CD8-T cell responses which has been shown to last for decades. Therefore, vaccine designs for emerging respiratory viruses need consideration and rational inclusion of CD8 epitopes to confer long term resistance (14).

This study demonstrates the existence of strong CD8 T-cell activating epitopes in the spike antigen and uncovers robust pre-existing CD8 T-cell immunity in unexposed donors. Several studies have reported pre-existing T-cell immunity in unexposed donors and attributed these to infections by common cold-causing human coronaviruses (7, 8, 10). Other studies, on the contrary, have reported a lack of pre-existing T-cell immunity in unexposed donors (15, 16). These differences can arise from the composition of peptide pools used in the assay since each group employed different selection strategies, the number of epitopes used by different groups was variable, differences among donor HLAs, dominant V and J genes in donors and the assay method. By using a smaller number of epitopes, and donors from two different regions of the globe, the USA and India, our findings confirm the existence of robust T-cell immunity in unexposed donors. Our results differ from other studies in two important aspects. First, published studies thus far have reported robust CD4 T-cell responses and a relatively weaker CD8 T-cell response in both unexposed and convalescent subjects, whereas we show strong CD8 T-cell response in both unexposed donors and convalescent patients. In fact, in our assays the IFN-γ response was significantly higher at 48 h than reported in other studies. Although, the assay conditions – such as the use of 15-mer peptides and the combination of cytokines used in the published studies could have favored a CD4 response over a CD8 response (8), however, we suspect that the epitope selection strategies and the use of large peptide pools by other groups may have masked the detection of strong CD8 T-cell epitopes. A second novel aspect of our study is the demonstration that the selected CD8 T-cell epitopes engaged cross-reactive TCRs in unexposed donors to mount a strong T-cell response. This finding has significant implications in COVID-19 vaccine development efforts (17) and the spread of the infection in different regions of the world (18).

To identify strong CD8 T-cell epitopes, we developed a novel TCR-binding algorithm OncoPeptVAC that selects epitopes favorable for TCR-binding. In all epitope screening methods, epitope selection is primarily based on class-I and II HLA-binding affinity, which predicts surface presentation of antigen in complex with HLA (19), but not the interaction of the peptide-HLA complex with a TCR (20). By incorporating features that predict TCR-binding of a peptide, our algorithm OncoPeptVAC successfully identified many CD8 T-cell epitopes in a small pool of 11 peptides used in T-cell activation assays. The TCR-binding algorithm is especially suitable for reducing the number of epitopes that need to be screened to identify robust CD8 T-cell activating epitopes. For example, our algorithm predicted 83 peptides from all SARS-CoV-2 proteins excluding ORF1, which is a much smaller number compared to the number of peptides screened in some of the published studies to identify pre-existing T-cell immunity (7, 8, 15, 21). A second factor that may have resulted in the identification of strong CD8 T-cell activating epitopes is the avoidance of epitope competition. Using a large pool of peptides to screen for T-cell responses ensures broad coverage of all HLAs, but has the disadvantage that strong immunogenic epitopes are not detected efficiently. Some of the peptides predicted by our algorithm produced >5% CD8 T-cell response in healthy donors by 14-days. In the same donors, the response from spike-S1 and S2 peptide pools containing 157 and 158 peptides respectively was much weaker. A similar finding was reported by Mateus et al. where deconvolution of peptide pools identified a single peptide that evoked 5-fold higher T-cell response compared to the pool (8). Also, important to note, that the strategy of using 15-mer peptides with overlapping 10 or 11-mer sequences may not identify immunodominant epitopes. For example, out of the 11-peptides tested in our assay, only three peptides were present in the spike peptide pools.

By using a smaller pool of immunodominant CD8 T-cell epitope, our study uncovered a fundamental feature of the host immune response to SARS-CoV-2 – the existence of cross-reactive TCRs to viruses, such as CMV and Influenza that recognizes SARS-CoV-2 antigen. An early and robust T cell response is driven by the size and the diversity of the TCR repertoire to a given antigen (22). The Pep-7 epitope derived from the RBD domain of SARS-CoV-2 spike antigen lacking homology to other coronaviruses expanded multiple public CDR3s recognizing immunodominant CMV epitope NLVPMVATV and Influenza epitope GILGFVFTL. Further, TCR analysis demonstrated that although a donor’s TCR repertoire contains many CMV-epitope-specific CDR3s, only a few are expanded in the presence of the SARS-CoV-2 peptide. For example, donor D225 TCR repertoire has 159 CMV and 249 Influenza CDR3s of which three and one were expanded respectively. Similarly, donor 089 carries 103 NLVPMVATV specific CDR3 of which only two expanded. These findings suggested the specificity of interaction between cross-reactive CDR3s and specific peptides from SARS-CoV-2. Significantly, the expanded CDR3s in the two donors D089 and D225 were different, even though they recognized the same CMV peptides. It has been documented that conserved features within CDR3-β allow recognition of the same pHLA complex within a group of diverse CDR3s (23). A robust antigen-specific T cell response utilizes a broad range of TCRs and for many viral infections, TCR usage diversity has been positively linked to disease outcomes (24-26). A diverse repertoire not only allows increased structural capacity to recognize variant epitopes (27) but increases the chances that high-affinity TCRs may be present in an individual (28). A recent large-scale study mapped a few immunogenic regions in the SARS-CoV-2 proteome responsible for expanding many unique TCRs in a large number of convalescent COVID-19 patients and unexposed healthy donors (21). Immunodominant epitopes reported in our study cover some of the “hotspot” regions identified by this large-scale study (21). Efforts to identify cross-reactive TCRs recognizing different antigens from diverse infectious organisms can lead to the development of broad-spectrum TCR-based therapeutics against infectious diseases.

We compared the All-peptide mix with spike-1 and 2 peptide pools on a small number of convalescent patients and identified a slightly higher CD8/IFN-γ response by the peptide mix in mild to moderate disease, compared to patients with asymptomatic or severe disease. Many studies have indicated that short and long-term protection against respiratory viruses requires CD8 T-cell immunity and antibody response alone is not sufficient (6, 29). In line with this observation, low plasma titers of neutralizing antibodies are detected in a large fraction of convalescent patients suggesting additional immune protective mechanisms, besides viral neutralization (30). On the contrary, high levels of neutralizing antibodies were associated with severe disease and ICU visits in many COVID-19 patients suggesting an imbalanced CD4 T-cell response is not optimal for protection (31-33). It has been challenging to demonstrate a strong CD8 T-cell response in COVID-19 patients in many studies. However, our findings along with a recent report from Peng et al. showed that a higher CD8 T-cell response correlated with a mild disease compared to patients with severe disease (15).

In conclusion, our study demonstrates strong pre-existing CD8 T-cell immunity in many unexposed donors contributed by the engagement of cross-reactive TCRs against common CMV and flu antigens. The presence of high-quality cross-reactive TCRs can protect individuals by mounting an early CD8 T-cell response and clearing the virus. Identifying additional immunodominant epitopes in SARS-CoV-2 and their cognate TCRs can become a powerful immune monitoring tool for assessing protective immunity against SARS-CoV-2 in the population.

## Supplementary Figures

**Figure S1. A-E**. Kinetics and magnitude of IFN-γ expression by CD8 T-cells in the presence of individual peptides from the 11-peptide-mix in unexposed donors.

**Figure S2. A-F**. Kinetics and magnitude of 4-1BB expression by CD8 T-cells in the presence of individual peptides from the 11-peptide-mix in unexposed donors.

**Figure S3. A-J**. Kinetics and magnitude of IFN-γ and 4-1BB expression by CD4 T-cells in the presence of 11-peptide-mix or individual peptides in unexposed donors.

**Figure S4. A-D**. Expression of T-cell activation and phenotype markers in different cell clusters from single-cell sequencing.

**Figure S5. A-D**. Expression of T-cell activation and phenotype markers in top-30 clonotypes from single-cell sequencing.

**Figure S6**. QC of cells from single cell transcriptomic analysis. The cells outside the red bars were excluded from the analysis (see Methods for details)

**Figure S7**. Clustering of cells without and with the expression of G1, G2/M and S cell cycle genes. Cell-type-specific clustering was achieved by regressing out the expression of cell cycle genes.

## METHODS

### T-cell epitope prediction

#### Dataset

Data on 371,865 T cell assays was collected from the IEDB (1). There were 105,673 CD8 T cell assays in total with 61,968 CD8 T cell assays with humans as a host. The CD8 T cell assays with HLA allele names and peptide lengths ranging 8-14 residues were further selected. HLA supertypes were replaced with their representative allele names, for example, HLA-A2 was replaced with HLA-A*02:01. The immunogenic peptide-HLA pairs tested on at least three donors with 100% response frequencies or at least tested on 5 donors with greater than 50% response frequencies were labelled as positive. The non-immunogenic peptide-HLA pairs tested on at least 3 donors with 0% response frequency were labelled as negative.

The final dataset contained 8,870 unique peptide-HLA pairs which were split randomly into 80% training and 20% test datasets. The training dataset had 884 immunogenic and 6,212 non-immunogenic peptide HLA pairs. The test dataset had 238 and 1,536 immunogenic and non-immunogenic peptide-HLA pairs, respectively.

#### Model

A deep Convolutional Neural Network (CNN) was implemented to predict immunogenicity of the peptide-HLA pair (provisional patent pending). The HLA alleles were represented as pseudo-sequences described as 34 amino acid residues (2). The peptide and HLA pseudo-sequences were converted to the two-dimensional (2D) feature matrices of 14×20 and 34×20 dimensions using BLOSUM encoding (3) respectively. Peptide sequences shorter than 14 residues were padded by zeroes to maintain 14×20 feature matrix dimensions. Peptide sequences were also encoded into 1×14 feature vector using the Kyte-Doolittle hydrophobicity scale (4). The HLA binding percentile ranks, and scores for each peptide-HLA pair were obtained using NetMHCpan-4.1 (5) and were appended to the Kyte-Doolittle hydrophobicity scale feature vector.

The peptide and HLA feature vectors were each processed by multiple 2D convolutional filters of two different sizes followed by max-pooling layers of the same sizes serially. The peptide and HLA max-pooled layers were concatenated and processed again with multiple 2D convolutional filters followed max-pooling layers. The 2 max-pooled layers were flattened-concatenated and then connected to a dense layer. The output of the peptide and HLA dense layer was concatenated with the hydrophobicity and HLA binding feature vector and again connected to two dense layers. The final output of dense layer was connected to the output neuron.

Three different versions of the CNN models were trained to evaluate if the hydrophobicity scale and HLA binding scores added to the performance of BLOSUM encoding. The first version, called OncoPeptVAC-2.0, was trained only using the BLOSUM encoding. The second version, called OncoPeptVAC-2.1, was trained using BLOSUM encoding and hydrophobicity indices. Final model version, called OncoPeptVAC-2.2, was trained using all 3 features, namely BLOSUM encoding, hydrophobicity indices and HLA binding scores. The hyperparameters of each model version were tuned based on model performance on the blind test dataset. The CNN was trained using 5-fold cross validation with the training dataset exclusively. The test dataset was solely used for model performance evaluation. Model performance was evaluated using AUC (area under ROC Curve) where an AUC of 0.5 represents random predictions and AUC of 1.0 represents the perfect predictions. The tensorflow library from Python programming language was used to implement the models.

#### Homology analysis

Full length shortlisted peptide sequences of SARS-CoV-2 were blasted against the spike proteins of other coronaviruses, OC43, NL63, 229E and HKU1. An E-value cutoff of 0.01 was used with a minimum cutoff of 11 amino acid residues was used to identify homologous peptides.

#### T cell activation assay

Unexposed donor PBMCs were obtained from the US and India for this study. PBMCs from the US were collected between 2016 - 2018 and purchased from Stemcell Technologies, Canada. PBMCs from India were collected between 2015 – 2018. COVID-19 convalescent patient blood from the US was purchased from PPA Research (USA) and the Indian samples were collected through hospitals. All participants in this study provided informed consent in accordance with protocols approved by the institutional review board. PBMCs were thawed, counted and analyzed using the diagnostic panel of antibodies (Table S5). PBMCs were rested overnight in RPMI containing 10% human serum (Table S5). For T-cell activation assays, 750,000 PBMCs were incubated either with DMSO (negative control) or with different peptide pools in 0.5 ml RPMI (Gibco) +10% Human AB serum (Sigma) + 10 ng/ml IL-15 and 10 IU of IL-2 (Stemcell Technologies, Canada). The culture media was replenished every three days with fresh media containing 10 IU of IL-2 and 10 ng/ml IL-15. On days 7, 14 and 21 of incubation, fresh peptides were added to the culture. For intracellular cytokine staining, cells were treated with Brefeldin A (BD Biosciences) for 5 hours, fixed and permeabilized using BD Lysis solution and Perm2 solutions respectively followed by staining with T-cell activation panel of antibodies (Table S5). Stained cells were analyzed in BD Accuri C6 Plus to detect the expression of activation markers IFN-γ and CD137 (4-1BB) on CD4 and CD8 T cells. Data was analyzed using BD Accuri C6 Plus software.

#### TCR sequencing and data analysis

200,000 PBMCS was removed after 48h, 7, 14 and 21 days from the T-cell activation assay and processed for bulk TCR sequencing.

#### Bulk TCR sequencing

TCR repertoire profiling was performed using the SMARTer TCR α/β Profiling Kit (Takara Bio, USA) according to the manufacturer’s protocol. RNA was isolated using the Qiagen RNA isolation kit. 10ng RNA from antigen-induced PBMCs were used as the starting material. The kit uses SMART technology (<u>S</u>witching <u>M</u>echanism <u>A</u>t 5’end of <u>R</u>NA <u>T</u>emplate) with 5’RACE to capture the entire V(D)J variable regions of TCR transcripts followed by two rounds of semi-nested PCR to obtain TCR-α and the β-chain. Libraries are prepared analyzed for quality and quantity. Sequencing is performed using the 2*300 MiSeq Reagent Kit v3 (Illumina, Inc.).

#### Single cell TCR sequencing and transcriptome profiling

For each sample, raw gene expression matrices were generated by Cell Ranger(v.3.0.2) coupled to the human reference version GRCh38. The gene expression data was analyzed by R software (v.3.4.4) with the Seurat package (2.3.4). In brief, Low-quality cells were removed if they met one of the following criteria: >75,000 unique molecular identifiers (UMIs); <500 or >7,500 genes; >10% UMIs derived from the mitochondrial genome; and >50% of transcripts contributed by top 50 genes (Figure S7). After removing low-quality cells, gene expression matrices were normalized by the NormalizeData function and features with high cell-to-cell variation were calculated using the FindVariableGenes function. Next, the expression of the S-phase and the G2-M phase genes were used to calculate the cell cycle score for all the cells using the CellCycleScoring feature. To generate unbiased clustering of cells ScaleData function was used, regressing out the expression of cell cycle genes, mitochondrial % and number UMI from the features (Figure S8). The dimensionality of the datasets was reduced by RunPCA function using variable features identified by FindVariableGenes function on linear-transformation scaled data generated by the ScaleData function. Next, the ElbowPlot and DimHeatmap functions were used to identify the true dimensionality of each dataset. Finally, cells were clustered using the FindClusters function and nonlinear dimensional reduction with the RunUMAP function using the Euclidean Distance feature. All details regarding the Seurat analyses performed in this work can be found in the website tutorial (https://satijalab.org/seurat/v2.4/pbmc3k_tutorial.html).

#### Cluster annotation and differential expression of genes

After nonlinear dimensional reduction and projection of all cells into two-dimensional space by UMAP, cells were clustered together according to common features. The FindAllMarkers function in Seurat was used to find markers for each of the identified clusters. Clusters were then classified and annotated based on expressions of canonical markers of particular cell types. Differential gene expression was performed using the FindAllMarkers function in Seurat with default parameters. We selected top-25 upregulated DEGs with maximum FDR value of 0.01 and annotated the clusters based on the expression of these upregulated genes. Next, we used MergeSeuratFunction to integrate the four datasets from the four treatment conditions and performed the steps of data normalization, feature extraction, regressing out features as described above. The heatmap and dot plots were generated using the DoHeatmap and DotPlot function in Seurat.

#### TCR V(D)J sequencing and analysis

Full-length TCR V(D)J segments were enriched using a Chromium Single-Cell V(D)J Enrichment kit according to the manufacturer’s protocol (10X Genomics). Demultiplexing, gene quantification and TCR clonotype assignment were performed using Cell Ranger (v.3.0.2) vdj pipeline with GRCh38 as reference. TCR diversity metric, containing clonotype frequency and barcode information, was obtained. Cells with at least one productive TCR α-chain (TRA) and one productive TCR β-chain (TRB) were retained for further analysis. Each unique TRA(s)-TRB(s) pair of TRA-TRB was defined as a clonotype. The presence of identical clonotypes at least in two cells were considered to be clonal, and the number of cells containing the same TRA-TRB pairs defined clonal amplification of a clonotype. Using barcode information, TCR clonotypes were projected on UMAP and Dot plots. Public TCRs were mapped to the IEDB and VDJDB annotated databases using the TRB sequence.

