## Supplemental Material for "Immunodominant T-cell epitopes from the SARS-CoV-2 spike antigen reveal robust pre-existing T-cell immunity in unexposed individuals"

### Table-S1. Unexposed Donor characteristics – Related to Figure-2

| Sl. No. | Donor ID | Age | Gender | HLA class-I and class-II |
|---|---|---|---|---|
| 1 | 116 | 25 | M | A*01:01:01:01/A*68:01:01:02, B*07:06:01/B*15:25:01, C*07:02:01:01/C*07:26:01, DRB1*04:03:01/DRB1*15:01:01:01, DQB1*03:02:01:01/DQB1*06:01:01 |
| 2 | 118 | 30 | M | A*24:02:01:01/A*24:02:01:01, B*07:06:01/B*39:01:01:03, C*07:02:01:01/C*12:04:02, DRB1*01:01:01/DRB1*09:01:02, DQB1*03:03:02:02/DQB1*05:01:01:02 |
| 3 | 122 | 28 | M | A*01:01:01:01/A*33:03:01, B*35:03:01:01/B*52:01:01:01, C*04:01:01:01/C*12:02:02:01, DRB1*14:04:01/DRB1*14:04:01, DQB1*05:03:01:01/DQB1*05:03:01:01 |
| 4 | 132 | 36 | M | A*01:01:01:01/A*31:01:02:01, B*35:03:01:01/B*51:01:01:01, C*04:01:01:01/C*14:07:01, DRB1*13:01:01:01/DRB1*14:07:01, DQB1*05:03:01:01/DQB1*06:03:01 |
| 5 | 142 | 27 | M | A*01:01:01:01/A*01:01:01:01, B*57:01:01/B*57:01:01, C*06:02:01:01/C*06:02:01:01, DRB1*07:01:01:01/DRB1*07:01:01:01, DQB1*03:03:02:01/DQB1*03:03:02:01 |
| 6 | 1610 | 39 | M | A*02:131/A*68:01:02:02, B*07:06:01/B*40:06:01:02, C*07:02:01:01/C*15:07, DRB1*04:03:01/DRB1*15:01:01:01, DQB1*03:02:01:01/DQB1*06:01:01 |
| 7 | 1615 | 41 | M | A*01:01:01:01/A*26:01:01:01, B*52:01:01:01/B*55:01:01, C*01:02:01:01/C*12:02:02:01, DRB1*04:110/DRB1*14:04:01, DQB1*03:02:01:01/DQB1*05:03:01:01 |
| 8 | 167 | 35 | M | A*01:01:01:01, A*02:03:01, B*37:01:01:01/B*51:06:01, C*06:02:01:01/C*14:02:01:01, DRB1*10:01:01:01/DRB1*14:04:01, DQB1*05:01:01:05/DQB1*05:03:01:01 |
| 9 | 169 | 24 | M | A*33:03:01:01/A*33:03:01:01, B*58:01:01:01/B*58:01:01:01, C*03:02:02:01/C*03:02:02:01, DRB1*03:01:01:01/DRB1*13:02:01:01, DQB1*02:01:01/DQB1*06:09:01:01 |
| 10 | 176 | 27 | M | A*01:01:01:01/A*68:01:02:02, B*15:18:01:02/B*51:06:01, C*07:04:01:01/C*14:02:01:01, DRB1*04:01:01:02/DRB1*15:01:01:01, DQB1*03:02:01:01/DQB1*06:01:01 |
| 11 | 089 | 42 | F | A*02:01:01 A*29:02:01, B*40:01:02 B*45:01:01, C*03:04:01 C*06:02:01 DRB1*07:01:01 DRB1*13:02:01 DRB3*03:01:01 DRB4*01:01:01:01 DQB1*02:02:01 DQB1*06:04:01 DPB1*04:01:01 DPB1*17:01 |
| 12 | 102C | 23 | M | A*11:01 A*24:02 B*07:02 B*40:01 C*03:04 C*07:02 DRB1*13:02 DRB1*15:01DRB3*03:01 DRB5*01:01 DQB1*06:02 DQB1*06:04 DPB1*02:01 |
| 13 | 225 | 42 | M | A*02:01:01 A*30:01:01 B*15:03:01 B*35:01:01 C*04:01:01 C*04:01:01 DRB1*03:01:01 DRB1*13:02:01 DRB3*02:02:01 DRB3*03:01:01 DQB1*02:01:01 DQB1*06:09:01 DPB1*01:01:02 DPB1*04:01:01 |
| 14 | 242 | 46 | M | A*03:01 A*24:02 B*07:02 B*40:01 C*03:04 C*07:02 HLA class-II not available |
| 15 | 384 | 50 | M | A*01:01 A*02:01 B*08:01 B*44:02 C*07:01 C*07:04 HLA class-II not available |
| 16 | 801C | 23 | M | A*11:01 A*29:02 B*38:01 B*44:03 C*12:03 C*16:01 DRB1*04:07 DRB1*07:01 DRB4*01:01 DRB4*01:03 DQB1*02:02 DQB1*03:01 DPB1*04:01 DPB1*04:01 |
| 17 | 907C | 19 | M | A*02:01 A*02:01 B*38:01 B*51:01 C*12:03 C*14:02 DRB1*03:01 DRB1*13:02 DRB3*01:01 DRB3*03:01 DQB1*02:01 DQB1*06:04 DPB1*01:01 DPB1*04:01 |

### Table S2. Clonally amplified CDR3s detected in the samples (Excel spreadsheet)

**Table S3. The number of cells present in each immune cell cluster and their distribution in different cell types**. For each treatment, the total number of cells sequenced on the 10X platform is shown in parenthesis. The number of cells belonging to each immune cell type is given. T-cell cluster expressed CD3E and CD3G but did not express any markers of CD4 or CD8 T-cells (refer to Figure S3-B). 'Not defined' cluster could not be assigned to any specific cell type based on marker expression (Figure S3).

| Cluster | DMSO (3399) | Spike-S1 (4991) | Spike-S2 (3965) | Pep-7 (4905) |
|---|---|---|---|---|
| CD8 | 2048 | 2452 | 1523 | 3033 |
| CD4 | 252 | 634 | 1102 | |
| γ/δ | 996 | 1378 | 727 | 827 |
| NKT | 34 | 373 | 540 | 236 |
| T-cell | | 86 | | 647 |
| Not defined | 69 | 68 | 73 | 162 |

**Table-S4. Convalescent patient characteristics – Related to Figure 5**

| Sl. No. | Patient ID | Age | Gender | Diagnosis | Severity |
|---|---|---|---|---|---|
| 1 | 442347 | 27 | M | RT-PCR | Asymptomatic |
| 2 | 443831 | 28 | M | RT-PCR | Asymptomatic |
| 3 | 443832 | 23 | F | RT-PCR | Asymptomatic |
| 4 | 443833 | 23 | F | RT-PCR | Asymptomatic |
| 5 | 443835 | 23 | F | RT-PCR | Asymptomatic |
| 6 | 50851 | 46 | F | RT-PCR | Asymptomatic |
| 7 | 51854 | 53 | F | RT-PCR | Asymptomatic |
| 8 | 53285 | 32 | M | RT-PCR | Asymptomatic |
| 9 | 43684 | 23 | F | RT-PCR | Mild to Moderate |
| 10 | 43683 | 49 | F | RT-PCR | Mild to Moderate |
| 11 | 442348 | 25 | M | RT-PCR | Mild to Moderate |
| 12 | 446201 | 55 | F | RT-PCR | Mild to Moderate |
| 13 | 446203 | 57 | M | RT-PCR | Mild to Moderate |
| 14 | 446205 | 30 | M | RT-PCR | Mild to Moderate |
| 15 | 446206 | 61 | M | RT-PCR | Severe |
| 16 | 443840 | 52 | M | RT-PCR | Severe |
| 17 | 56034 | 53 | M | RT-PCR | Severe |
| 18 | 63185 | 56 | F | RT-PCR | Severe |
| 19 | 446202 | 53 | M | RT-PCR | Severe |

**Table S5. Cell Culture and FACS reagents.**

| Reagents used for culturing PBMCs | | | |
|---|---|---|---|
| **S.No** | **Reagent** | **Catalog No.** | **Supplier** |
| 1 | Gibco™ RPMI 1640 Medium | 11875085 | Gibco |
| 2 | Human Serum | H4522 | Sigma-Aldrich, USA |
| 3 | Gibco™ Penicillin-Streptomycin (10,000 U/mL) | 15-140-122 | Gibco |
| 4 | Recombinant Human IL-15 (10µg) | 200-15 | Stemcell Technologies |
| 5 | Human Recombinant IL-2 (10µg) | 78036.1 | Stemcell Technologies |

| Antibodies for T cell diagnostics & activation | | | |
|---|---|---|---|
| **PBMC Diagnostic Panel** | | | |
| **S.No** | **Antibody** | **Catalog No.** | **Supplier** |
| 1 | CD8 APC (Clone: SK1) | 17008742 | eBioscence |
| 2 | CD45RO FITC (Clone: UCHL1) | 11045742 | eBioscence |
| 3 | CD56 FITC (Clone: TULY56) | 11056642 | eBioscence |
| 4 | CD57 FITC (Clone: TB01) | 11057742 | eBioscence |
| 5 | CD14 PE (Clone: 61D3) | 12014942 | eBioscence |
| 6 | CD16 PE (Clone: CB16) | 12016842 | eBioscence |
| 7 | CD4 PerCP - C5.5 (Clone: OKT4) | 45004842 | eBioscence |
| 8 | TCR γδ FITC/ TCR γδ BV 421 (Clone: 11F2) | 347903/744870 | BD Biosciences |
| **T cell Activation panel** | | | |
| 9 | CD8 PerCP - C5.5 (Clone: SK1) | 344710 | Biolegend |
| 10 | CD4 V450/CD4 BV510 (Clone: SK3) | 651849/344634 | BD/Biolegend |
| 11 | IFN γ APC (Clone: 4S.B3) | 502512 | Biolegend |
| 12 | TCR γδ FITC/ TCR γδ PE-Cy™7 (Clone: 11F2) | 347903/655410 | BD Biosciences |
| 13 | 41BB PE / 41BB FITC (Clone: 4B4 (4B4-1) | 12-1379-42/11137942 | eBioscence |

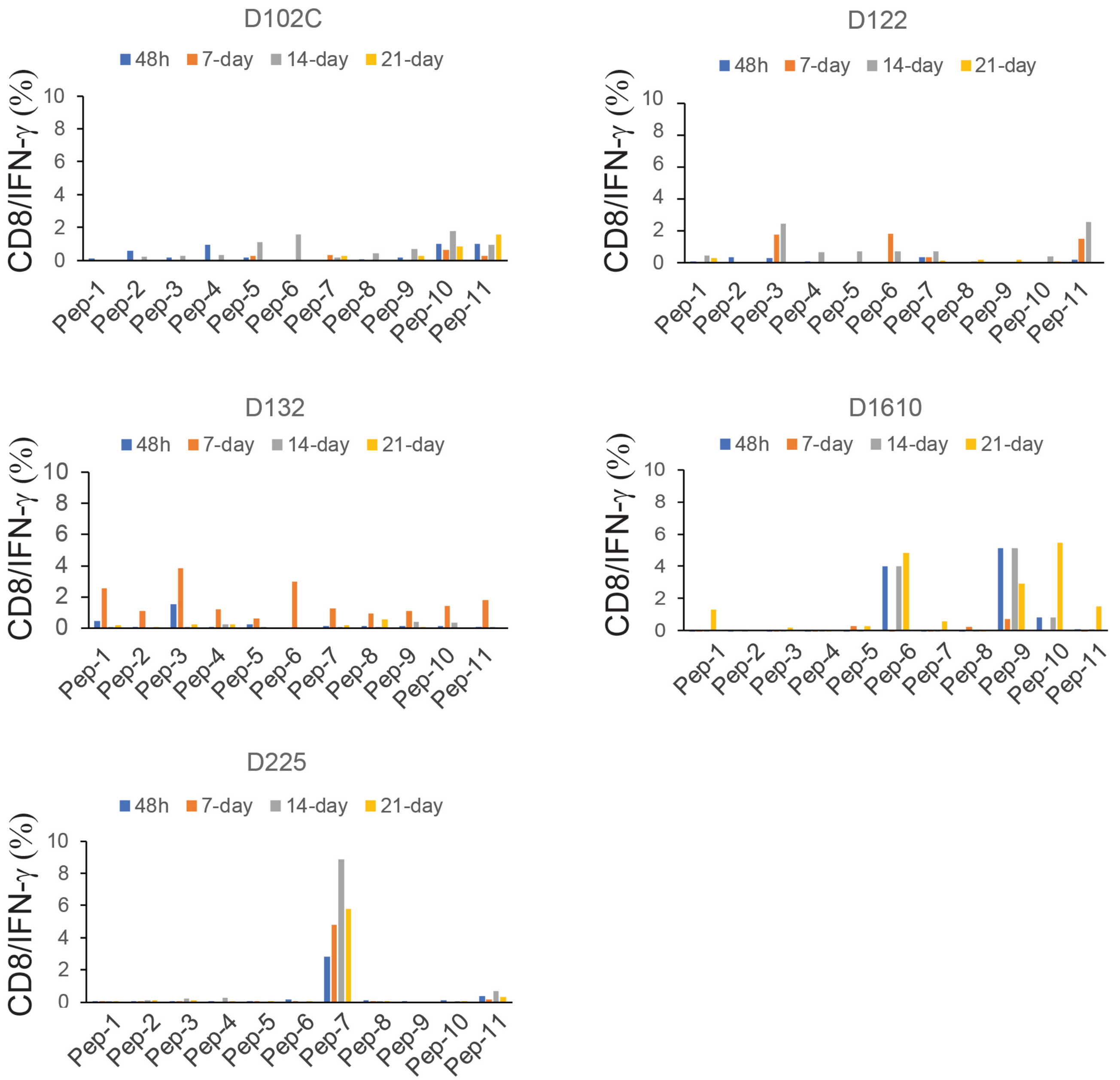
D102C
D122
D132
D1610
D225
48h
7-day
14-day
21-day
CD8/IFN-γ (%)
0
2
4
6
8
10
Pep-1
Pep-2
Pep-3
Pep-4
Pep-5
Pep-6
Pep-7
Pep-8
Pep-9
Pep-10
Pep-11

Figure S1

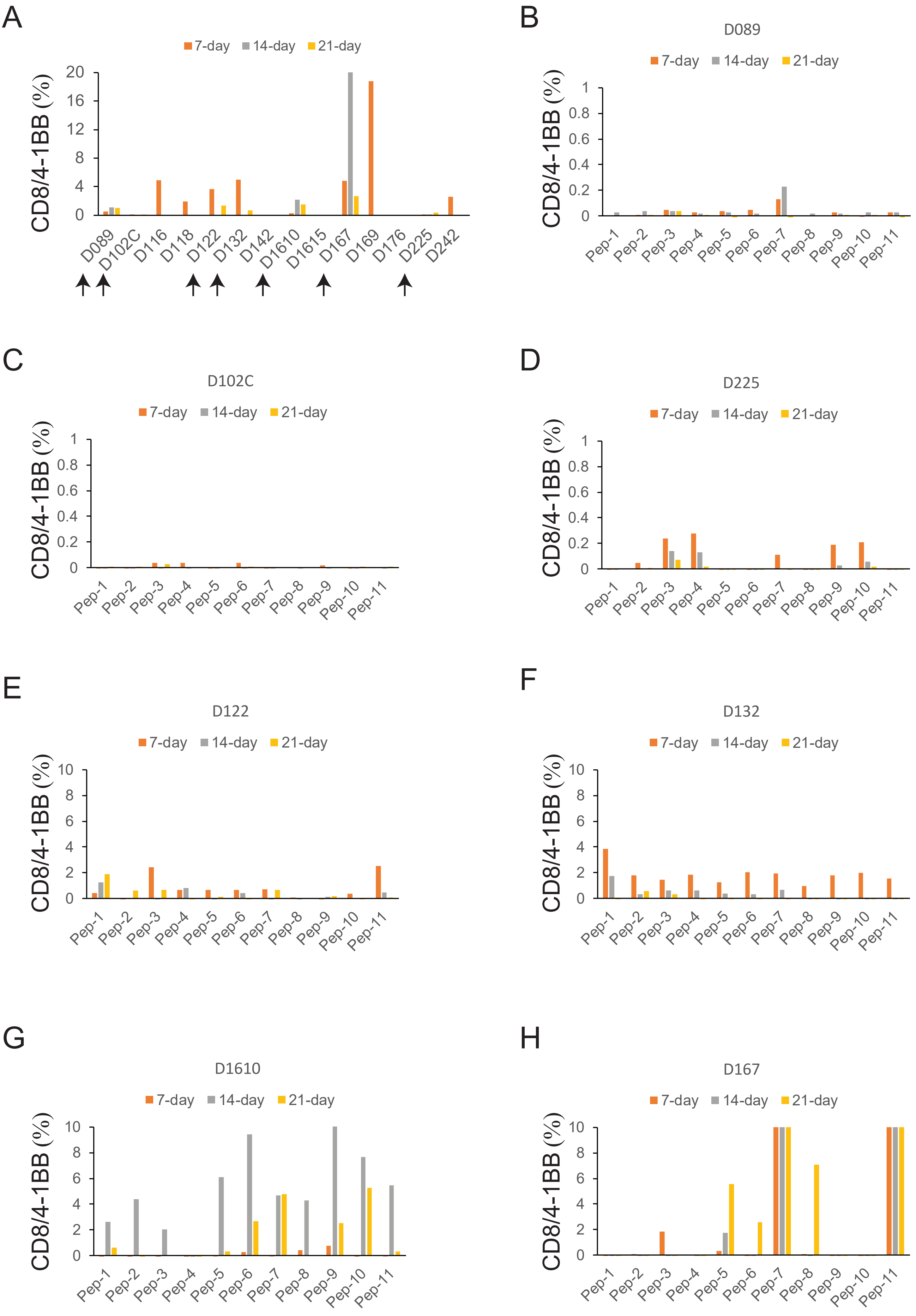
A
7-day 14-day 21-day
CD8/4-1BB (%)
D089 D102C D116 D118 D122 D132 D142 D1610 D1615 D167 D169 D176 D225 D242
B
D089
7-day 14-day 21-day
CD8/4-1BB (%)
Pep-1 Pep-2 Pep-3 Pep-4 Pep-5 Pep-6 Pep-7 Pep-8 Pep-9 Pep-10 Pep-11
C
D102C
7-day 14-day 21-day
CD8/4-1BB (%)
Pep-1 Pep-2 Pep-3 Pep-4 Pep-5 Pep-6 Pep-7 Pep-8 Pep-9 Pep-10 Pep-11
D
D225
7-day 14-day 21-day
CD8/4-1BB (%)
Pep-1 Pep-2 Pep-3 Pep-4 Pep-5 Pep-6 Pep-7 Pep-8 Pep-9 Pep-10 Pep-11
E
D122
7-day 14-day 21-day
CD8/4-1BB (%)
Pep-1 Pep-2 Pep-3 Pep-4 Pep-5 Pep-6 Pep-7 Pep-8 Pep-9 Pep-10 Pep-11
F
D132
7-day 14-day 21-day
CD8/4-1BB (%)
Pep-1 Pep-2 Pep-3 Pep-4 Pep-5 Pep-6 Pep-7 Pep-8 Pep-9 Pep-10 Pep-11
G
D1610
7-day 14-day 21-day
CD8/4-1BB (%)
Pep-1 Pep-2 Pep-3 Pep-4 Pep-5 Pep-6 Pep-7 Pep-8 Pep-9 Pep-10 Pep-11
H
D167
7-day 14-day 21-day
CD8/4-1BB (%)
Pep-1 Pep-2 Pep-3 Pep-4 Pep-5 Pep-6 Pep-7 Pep-8 Pep-9 Pep-10 Pep-11

Figure S2

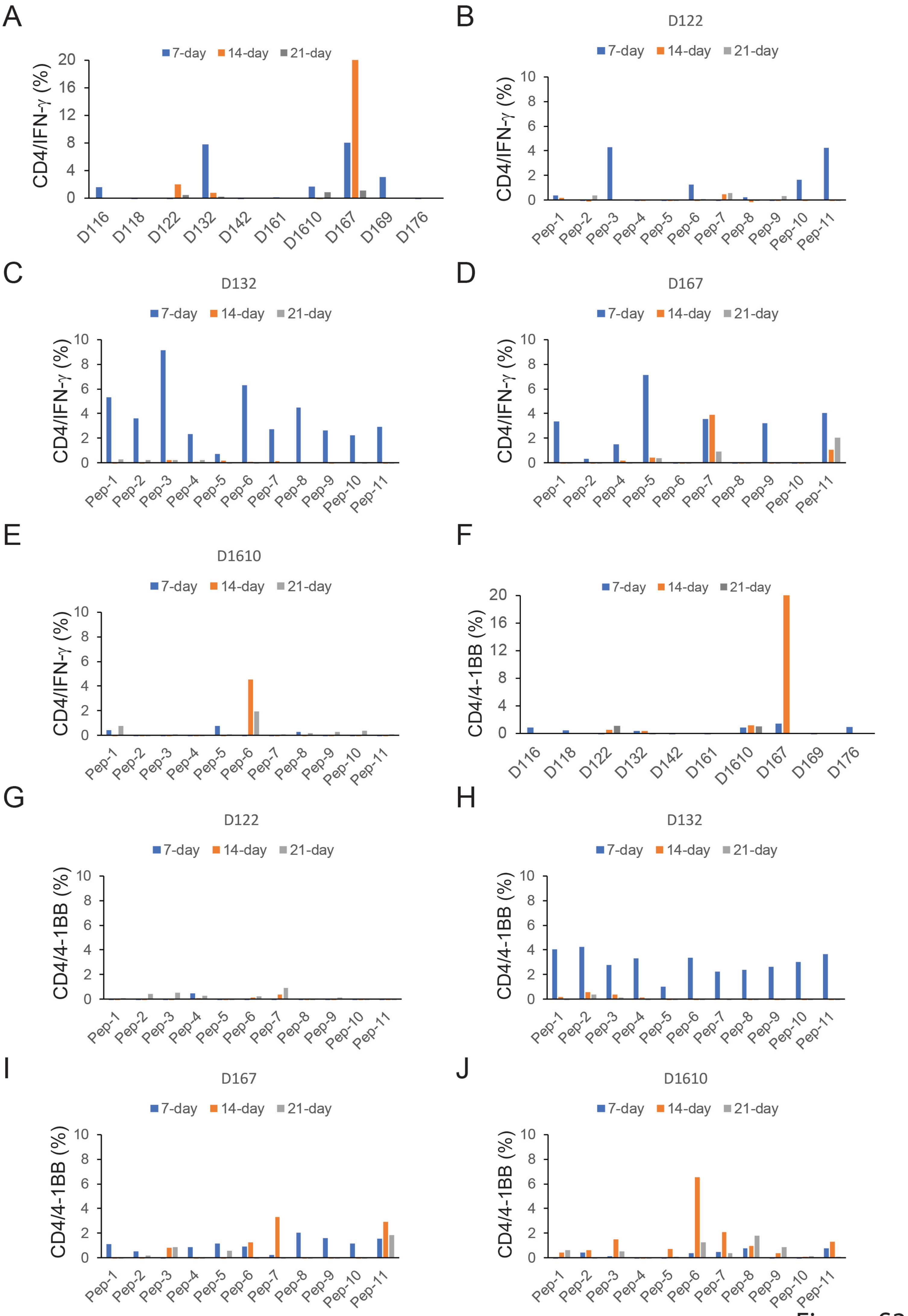
A
B
C
D
E
F
G
H
I
J
D122
D132
D167
D1610
D122
D132
D167
D1610
7-day
14-day
21-day
CD4/IFN-γ (%)
CD4/4-1BB (%)
D116
D118
D122
D132
D142
D161
D1610
D167
D169
D176
Pep-1
Pep-2
Pep-3
Pep-4
Pep-5
Pep-6
Pep-7
Pep-8
Pep-9
Pep-10
Pep-11

Figure S3

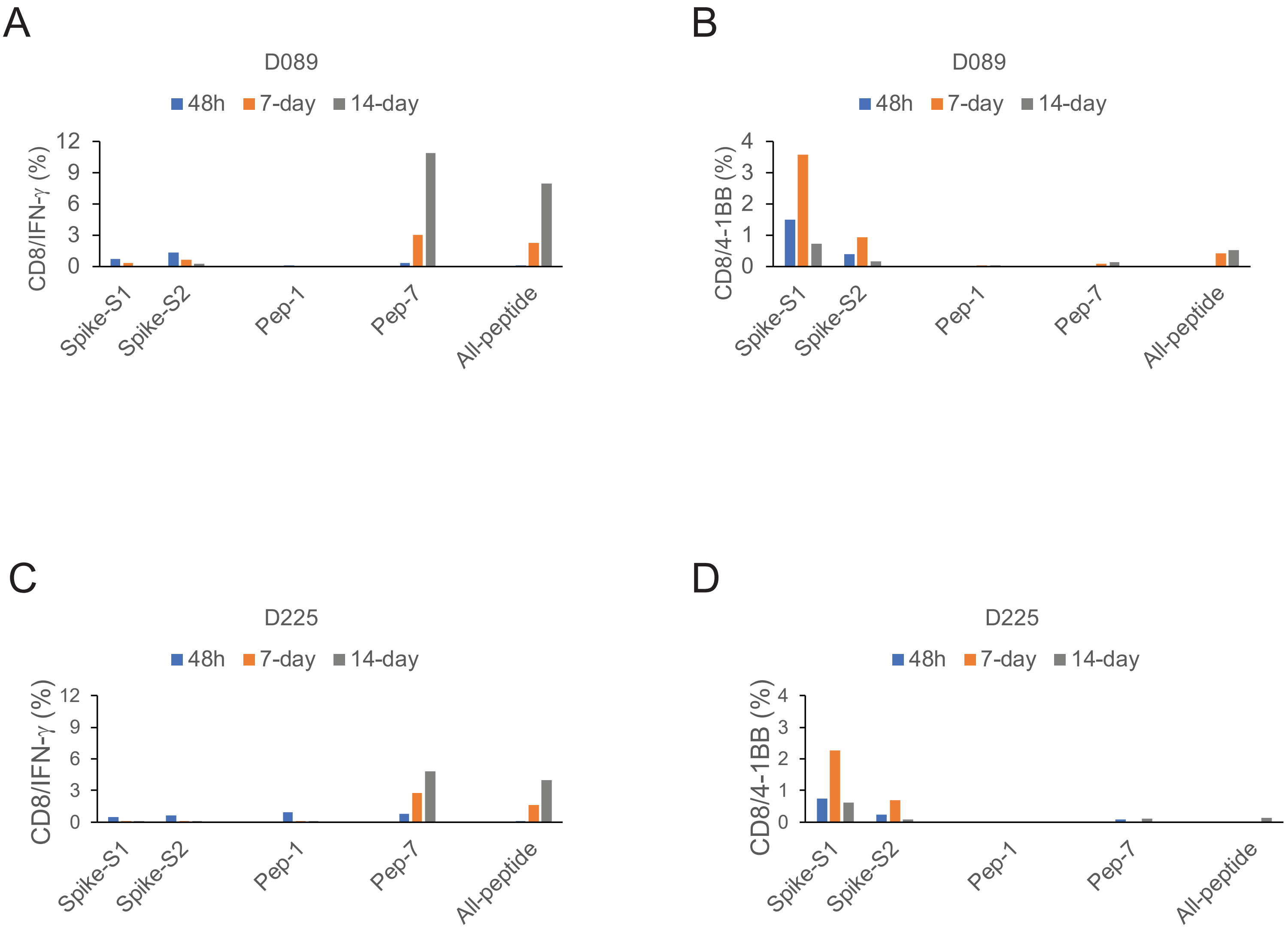
A
D089
48h
7-day
14-day
CD8/IFN-γ (%)
0
3
6
9
12
Spike-S1
Spike-S2
Pep-1
Pep-7
All-peptide
B
D089
48h
7-day
14-day
CD8/4-1BB (%)
0
1
2
3
4
Spike-S1
Spike-S2
Pep-1
Pep-7
All-peptide
C
D225
48h
7-day
14-day
CD8/IFN-γ (%)
0
3
6
9
12
Spike-S1
Spike-S2
Pep-1
Pep-7
All-peptide
D
D225
48h
7-day
14-day
CD8/4-1BB (%)
0
1
2
3
4
Spike-S1
Spike-S2
Pep-1
Pep-7
All-peptide

Figure S4

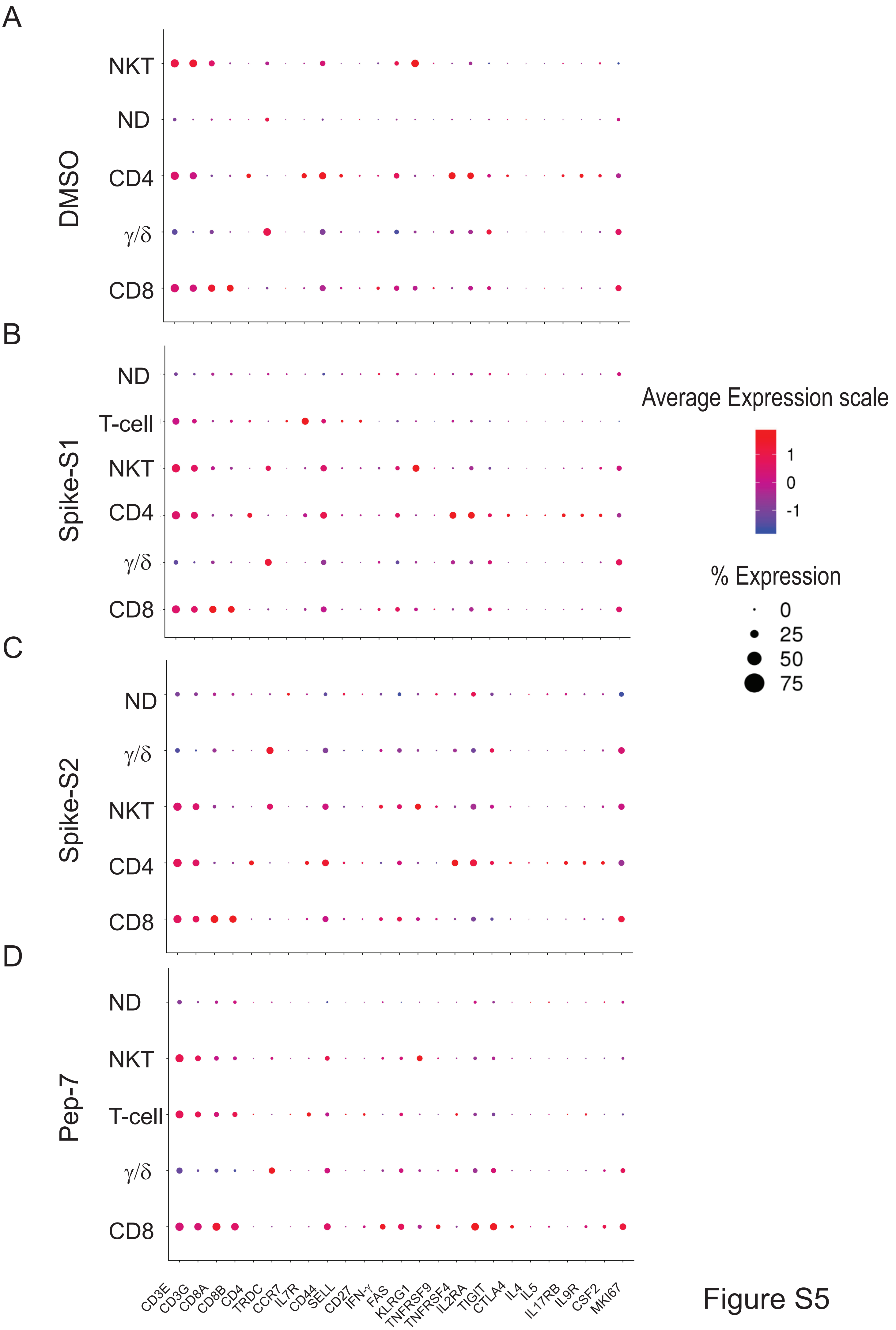
A
B
C
D
DMSO
Spike-S1
Spike-S2
Pep-7
NKT
ND
CD4
γ/δ
CD8
T-cell
Average Expression scale
1
0
-1
% Expression
0
25
50
75
CD3E
CD3G
CD8A
CD8B
CD4
TRDC
CCR7
IL7R
CD44
SELL
CD27
IFN-γ
FAS
KLRG1
TNFRSF9
TNFRSF4
IL2RA
TIGIT
CTLA4
IL4
IL5
IL17RB
IL9R
CSF2
MKI67

Figure S5

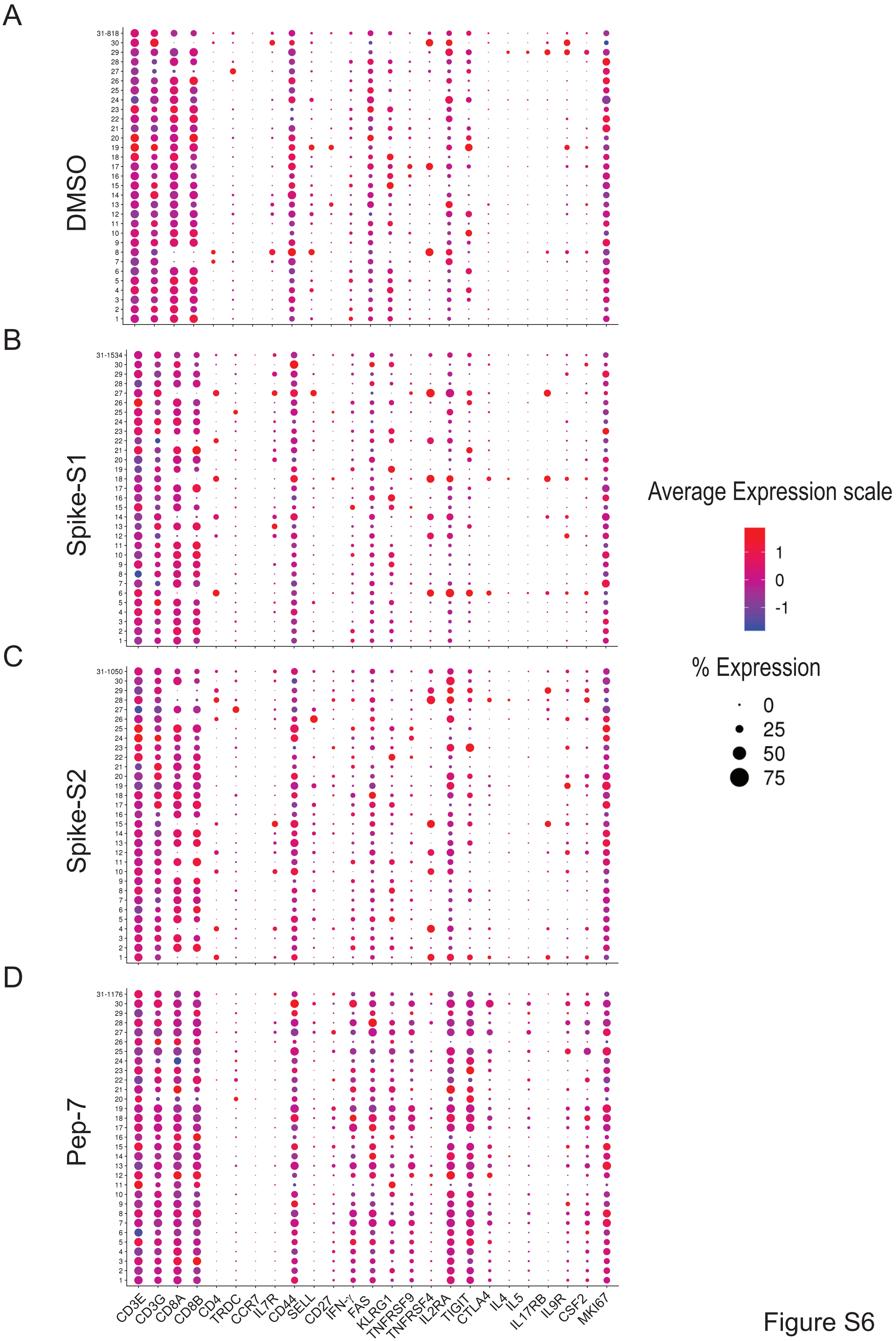
A
DMSO
B
Spike-S1
C
Spike-S2
D
Pep-7
Average Expression scale
1
0
-1
% Expression
0
25
50
75
CD3E
CD3G
CD8A
CD8B
CD4
TRDC
CCR7
IL7R
CD44
SELL
CD27
IFN-γ
FAS
KLRG1
TNFRSF9
TNFRSF4
IL2RA
TIGIT
CTLA4
IL4
IL5
IL17RB
IL9R
CSF2
MKI67

Figure S6

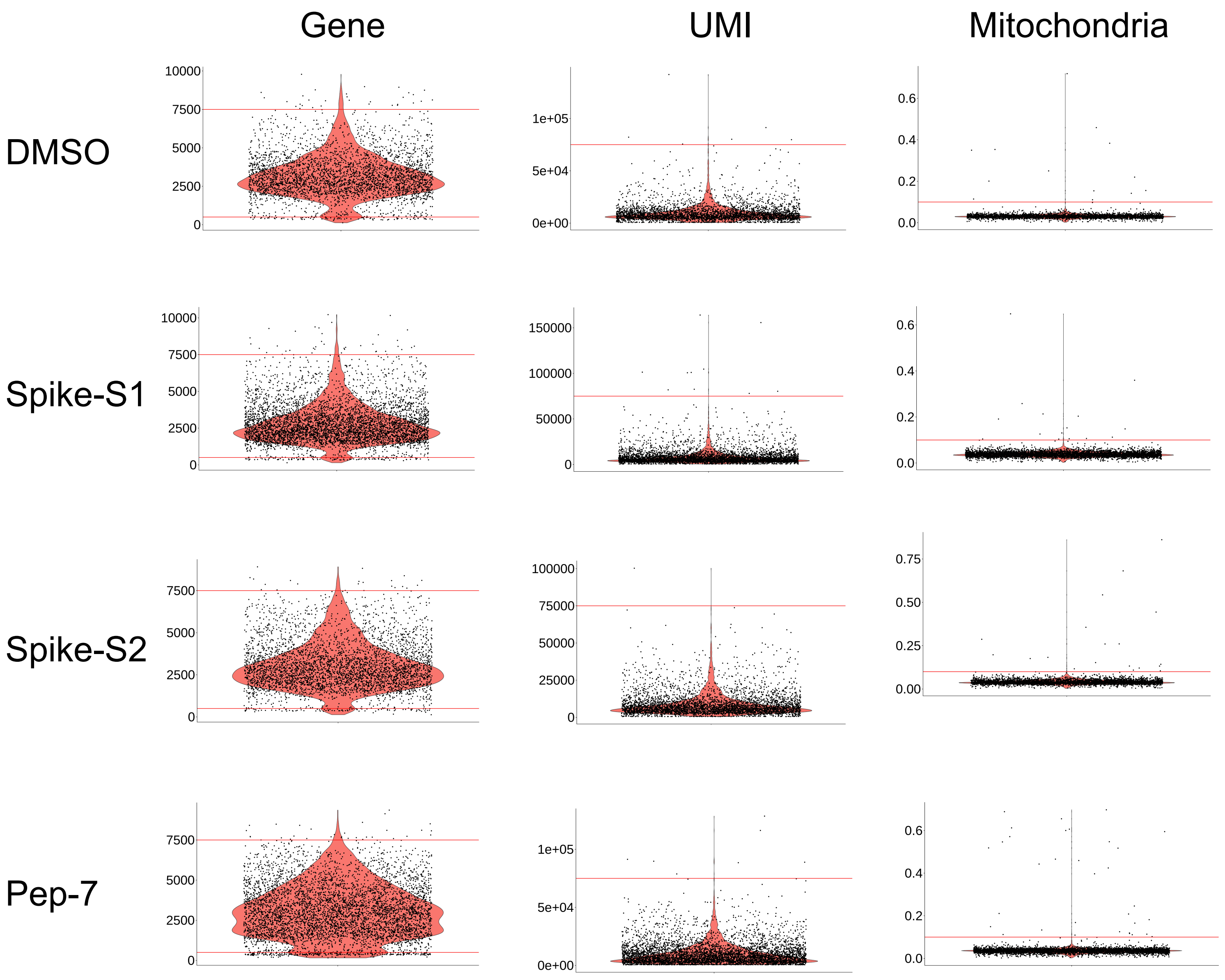

Gene
UMI
Mitochondria
DMSO
Spike-S1
Spike-S2
Pep-7
10000
7500
5000
2500
0
1e+05
5e+04
0e+00
150000
100000
50000
75000
25000
0.75
0.50
0.25
0.00
0.6
0.4
0.2
0.0


Figure S7

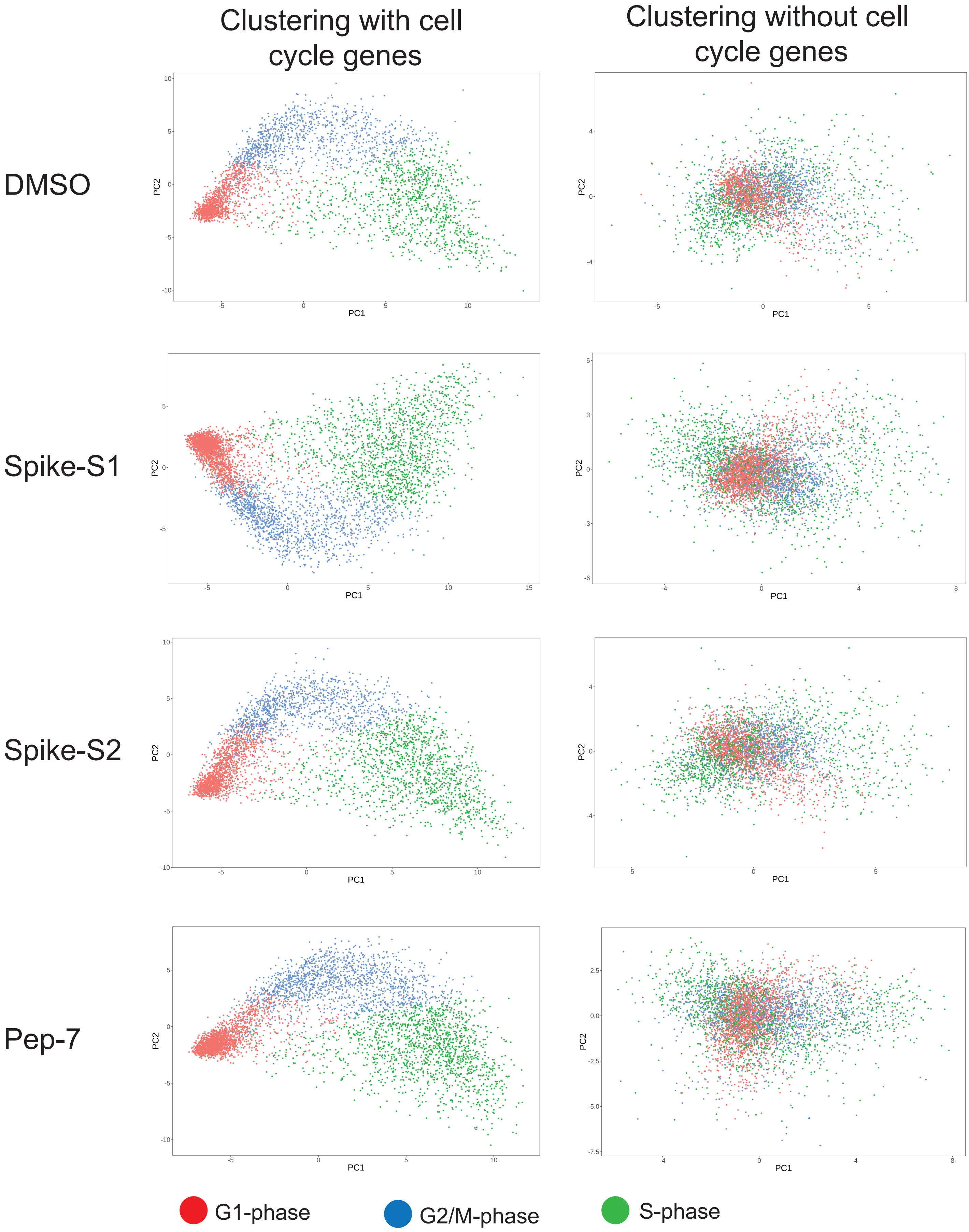
Clustering with cell cycle genes
Clustering without cell cycle genes
DMSO
Spike-S1
Spike-S2
Pep-7
PC1
PC2
G1-phase
G2/M-phase
S-phase

Figure S8
